# A sensory cell diversifies its output by varying Ca^2+^ influx-release coupling among presynaptic active zones for wide range intensity coding

**DOI:** 10.1101/2020.06.06.137919

**Authors:** Özge Demet Özçete, Tobias Moser

## Abstract

The cochlea encodes sound intensities ranging over six orders of magnitude which is collectively achieved by functionally diverse spiral ganglion neurons (SGNs). However, the mechanisms enabling the SGNs to cover specific fractions of the audible intensity range remain elusive. Here we tested the hypothesis that intensity information, fully contained in the receptor potential of the presynaptic inner hair cell (IHC), is fractionated via heterogeneous synapses. We studied the transfer function of individual active zones (AZs) using dual-color Rhod-FF and iGluSnFR imaging of Ca^2+^ and glutamate release. AZs differed in the voltage dependence of release: AZs residing at the IHCs’ pillar (abneural) side activate at more hyperpolarized potentials and typically showed tight control of release by few Ca^2+^-channels. We conclude that heterogeneity of voltage dependence and release-site coupling of Ca^2+^-channels among the AZs varies synaptic transfer within individual IHCs and, thereby, likely contributes to the functional diversity of SGNs.

## Introduction

Neural systems employ functional diversity to achieve the complexity of behavior. Diversity is implemented at several levels, i.e. circuits, neurons, and subcellular functional units such as synapses. For instance, the auditory system copes with the task of encoding a wide range of sound intensities by employing two types of sensory cells: outer hair cells to actively compress the range of mechanical inputs and IHCs to adaptively encode at synapses with primary auditory neurons (type I SGNs). Based on their physiology, these neurons can be classified into three functional subtypes, namely low, medium, and high spontaneous rate (SR) SGNs differing in the thresholds and dynamic range of sound encoding^1–5^. This functional SGN diversity appears at all tonotopic places of the cochlea and the subtypes can even innervate the same IHC^6^.

Functional diversity of SGNs relates to the heterogeneity of their molecular profile, morphology, afferent and efferent synaptic connectivity. In the cat^6^, back-tracing experiments linked morphology to function and showed that low-SR SGNs have thinner radial fibers (peripheral neurites) with fewer mitochondria than high-SR. Low-SR SGNs preferentially innervate the modiolar (or neural) side of the IHC, where they face larger and more complex presynaptic AZs^6–9^. Larger and more complex AZs at the modiolar side of IHCs were also found in mouse^10–12^, guinea pig^13,14^, and gerbil^15^, suggesting some features of SGN morphology and synaptic connectivity may be shared across species. In the mouse, RNA sequencing of individual SGNs indicated three distinct subtypes^16–18^ that were suggested to correspond to low, medium, and high-SR SGNs based on the spatial segregation of their IHC innervation.

In mouse IHCs, AZ size correlates with the number of Ca^2+^-channels (approximately 30 to 300)^19^ and consequently with the maximal Ca^2+^-influx at the AZ^11,19,20^. Should the analogy to the innervation pattern in the cat cochlea apply, it is odd that Ca^2+^-triggered glutamate release from modiolar AZs, with large size and Ca^2+^-influx, were to drive low-SR SGNs. A possible solution to this conundrum came from the finding that Ca^2+^-influx at the modiolar AZs requires stronger depolarization than at the pillar ones^11^. In other words, Ca^2+^-channels at modiolar AZs would be mostly closed at the IHC resting potential and require stronger receptor potentials to activate, which could explain the low SR and higher thresholds of their postsynaptic SGNs.

Whether and precisely how such heterogeneous properties of presynaptic Ca^2+^-signaling relate to glutamate release and, consequently to SGN firing remains to be elucidated. Exocytosis of readily releasable synaptic vesicles (SVs) in mature IHCs relates near-linearly to Ca^2+^ influx, when varying the number of open Ca^2+^-channels^21–23^. Given this near-linear dependence, one would assume that the heterogeneous Ca^2+^-signaling propagates into a concomitant diversity of transmitter release. However, this remains to be studied at the single-synapse level ideally for several AZs of a given IHC. In fact, the near-linear Ca^2+^-dependence of IHC exocytosis was proposed to result from summation of individual AZs with supralinear Ca^2+^-dependence but different operating points^24^. Moreover, a recent study of cerebellar synapses highlighted how differences in Ca^2+^ channel-release coupling diversify synaptic transfer^25^.

Here we studied the synaptic transfer function and underlying Ca^2+^-dependence of release at individual IHC-SGN synapses by combining IHC patch-clamp with imaging of synaptic Ca^2+^ influx and glutamate release. To detect glutamate release, we utilized the fluorescent glutamate reporter iGluSnFR^26^ that we targeted to the postsynaptic SGN bouton. Our results suggest that IHCs vary the voltage-dependence of Ca^2+^-channels as well as their control of release-sites among their AZs. This likely enables IHCs to signal the information contained in the receptor potential into complementary neural channels for encoding the entire audible range of sound.

## Results

### Optical detection of glutamate release at individual inner hair cell synapses

Fluorescence imaging allows analysis of individual IHC AZs^11,20,27^ due to their large nearest neighbor distance (∼2µm)^28^. To image glutamate release, we targeted iGluSnFR^26^ to the postsynaptic SGN membrane. We injected the round window of WT mice at postnatal day (P)5-7 with adeno-associated virus (AAV9, human synapsin promoter) to drive largely uniform SGN expression of iGluSnFR with several transduced afferent boutons per IHC (Fig. 1a).

**Figure 1:**
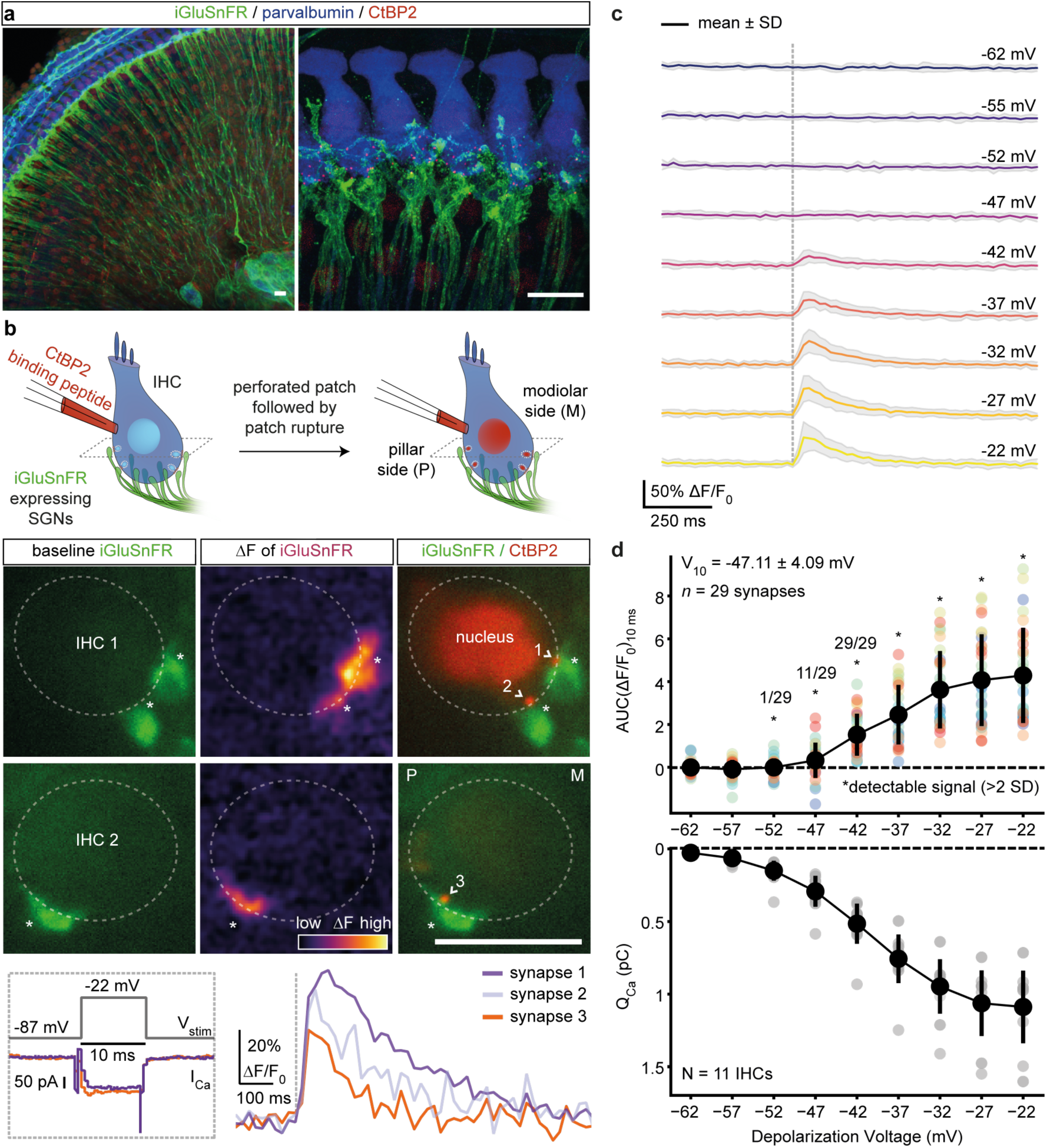
Optical detection of glutamate release at individual IHC synapses: low and variable voltage threshold. **a.** Maximum intensity projections of the organ of Corti from the right ear of a P17 mouse injected at P6 with 1-2 µl of AAV9.*hSyn*.iGluSnFR suspension, immunolabeled for iGluSnFR (GFP), IHCs, OHCs and SGNs (parvalbumin) and synaptic ribbons and nucleus (CtBP2/RIBEYE). Close-up (right panel) highlights IHCs innervated by several iGluSnFR-expressing SGN boutons. **b.** Simultaneous perforated patch-clamp and imaging of IHCs. IHCs were patch-clamped from the pillar side, with a patch pipette containing TAMRA-conjugated CtBP2-binding peptide. Towards the end of the recording, the membrane patch was ruptured to fill the IHC with the peptide, which stains synaptic ribbons and nucleus. (Left) Two exemplary confocal sections of IHCs showing baseline fluorescence of iGluSnFR-expressing afferent boutons from both modiolar (IHC1) and pillar (IHC2) side of the cell. Glutamate release from IHCs was evoked upon 10-ms-long step depolarizations to −22 mV from a holding potential of −87 mV (see bottom, 1.3 mM [Ca^2+^]_e_), and detected as fluorescence change (ΔF) of iGluSnFR signal (recorded at 50 Hz) located on the SGN membrane. (Right) Overlaid images of the IHC1 and IHC2, displaying the boutons (iGluSnFR) and the synaptic ribbons (CtBP2), after the recording. (*: transduced afferent boutons, >: synaptic ribbons) (Scale bars: 10 µm) **c.** Average ΔF/F_0_ iGluSnFR traces in response to 10-ms-long step depolarizations from the holding potential (−87 mV) to a voltage within the physiologically relevant range of receptor potentials: from −62 mV to −22 mV (applied in pseudo-randomized order, step-size 5 mV, perforated patch-clamp, 1.3 mM [Ca^2+^]_e_, *n* = 29 boutons, N = 11 IHCs from 9 mice). Shaded areas show ± SD. **d.** The voltage threshold of glutamate release was low and variable (−47.11 ± 4.09 mV, mean ± SD). The area under the curve of the iGluSnFR signal (top; AUC(ΔF/F_0_)_10ms_) from c and corresponding whole-cell Q_Ca_ (bottom) (mean ± SD). Detectable signals were defined here if the peak iGluSnFR signal was 2 times higher than baseline SD. All synapses had detectable signals in response to depolarizations ≥ −42 mV. (See also Supplementary Figs. 1-2)

Using apicocochlear organs of Corti, acutely dissected after the onset of hearing (P15-19), we patch-clamped IHCs and simultaneously imaged postsynaptic iGluSnFR fluorescence by spinning-disk confocal microscopy^11^. Fig.1b shows two exemplary IHCs innervated by iGluSnFR-expressing afferent boutons either at their modiolar (IHC1) or pillar (IHC2) side in the given confocal sections. Sizable changes in iGluSnFR fluorescence (ΔF-iGluSnFR) were evoked by brief (10-ms-long) step depolarizations to −22 mV (Fig. 1b). Toward the end of the perforated-patch recording, we ruptured the membrane patch and introduced a TAMRA-conjugated dimeric ribbon-binding peptide to identify individual AZs (Fig. 1b). When imaging, we purposely avoided the basal pole of the IHCs in which separating individual postsynaptic boutons is more challenging given the high synapse density^10,28^.

To probe the specificity of ΔF-iGluSnFR for reporting Ca^2+^-mediated glutamate release, we tested the effect of the Ca^2+^-channel blocker Zn^2+^. ΔF-iGluSnFR triggered by step depolarizations gradually decreased and partially recovered upon Zn^2+^ application and wash-out, respectively (Supplementary Fig. 1a-b). To assess potential adverse effects of iGluSnFR expression on auditory signaling, we recorded auditory brainstem responses (ABR) at P29 (∼23 days after the AAV-injection). ABR waveforms and thresholds of the injected and non-injected (control) ears were comparable (Supplementary Fig. 1c) despite efficient transduction and iGluSnFR expression of SGNs. In conclusion, AAV-mediated expression of iGluSnFR in SGNs is suitable for studying IHC glutamate release with high specificity and does not obviously alter auditory physiology.

### Deciphering synaptic glutamate release from IHC AZs

To compare IHC exocytosis on a single-synapse versus whole-cell level, we measured synaptic ΔF-iGluSnFR simultaneously with well-established whole-cell membrane capacitance changes (ΔC_m_)^29^. To assess the sensitivity and saturation of ΔF-iGluSnFR, we applied stimuli of different durations (2 ms to 100 ms, in pseudo-randomized order) in near-physiological conditions (perforated-patch configuration, 1.3 mM extracellular Ca^2+^ concentration ([Ca^2+^]_e_)) (Supplementary Fig. 1d). ΔF-iGluSnFR became significant at 2 ms (*p* < 0.0001), while ΔC_m_ were detectable only at 5 ms (*p* = 0.004, data not shown, See Methods). To evaluate a potential saturation of iGluSnFR by glutamate release, we related both the peak and area under the curve of the iGluSnFR signal (hereafter referred to as iGluSnFR-AUC) to the corresponding ΔC_m_. Both measures showed a positive correlation with ΔC_m_ (Pearson’s *r* = 0.63, *p* < 0.0001 and *r* = 0.66, *p* < 0.0001, Supplementary Fig. 1f-g), indicating their robust report of exocytosis for depolarizations up to at least 100 ms. Different from postsynaptic ΔF-iGluSnFR, ΔC_m_ also reports extrasynaptic exocytosis^22^, likely contributing to the sublinear relationship of both measures for longer stimuli.

Next, we compared SV pool dynamics by synaptic iGluSnFR-AUC versus whole-cell ΔC_m_. To estimate the dynamics of RRP and sustained exocytosis, we fitted the sum of an exponential and a linear function. The resulting time constants of RRP depletion were 11.39 ms for iGluSnFR-AUC and 7.94 ms for ΔC_m_ (Supplementary Fig. 1e, h, and see Methods). By estimating an average AZ RRP of ∼10 SVs from ΔC_m_-measurements, we obtained an average change in iGluSnFR-AUC (0.23 arbitrary units) per SV. The sustained component of exocytosis amounted to 42.7 a.u./s (∼185 SV/s per AZ) for the recorded synapses compared to 242 fF/s (504 SV/s per AZ) (Supplementary Fig. 1h). The faster rate derived from whole-cell likely reflects the contribution of extrasynaptic exocytosis^.22^. Hence, ΔF-iGluSnFR can detect smaller amounts of exocytosis than whole-cell ΔC_m_ and, on average, reports similar SV pool dynamics for single AZs.

Next, we studied how the RRP is recruited by brief graded depolarizations in the range of physiological receptor potentials^30^. Using near-physiological conditions (perforated-patch configuration, 1.3 mM [Ca^2+^]_e_), we applied 10-ms-long step depolarizations from the holding potential (−87 mV) to −62 mV in 5 mV increments up to −22 mV (applied in pseudo-randomized order, Fig. 1c-d; Supplementary Fig. 2a-b). We analyzed the voltage dependence of glutamate release by fitting Boltzmann functions to iGluSnFR-AUC (See Methods; Supplementary Fig. 2c-d). The voltage threshold (V_10,_ defined as potential eliciting 10% of the maximum response) for glutamate release on average was −47.11 mV with considerable variance (SD: 4.09 mV, Supplementary Fig. 2c-d). This is close to the threshold of activation of Ca_V_1.3 Ca^2+^-channels (∼-60 mV)^31^ and the *in vivo* resting membrane potential of IHCs (∼-55 mV)^32^. The variable voltage thresholds for glutamate release among the individual IHC synapses hints to differences in their transfer functions.

### Sequential dual-color imaging of Ca^2+^ signal and glutamate release at single AZs

How the opening of Ca_V_1.3 Ca^2+^-channels translates into glutamate release critically shapes synaptic transfer and is determined by the topography of Ca^2+-^channels and SV release-sites. Previous studies evaluating the summed activity of several AZs indicate that a few channels in nanoscale proximity govern the [Ca^2+^] driving fusion of individual SVs at mature hair cell synapses (Ca^2+^-nanodomain-like control of exocytosis)^21–23,33^. In the Ca^2+^-nanodomain-like control of release, a near-linear relation between release and Ca^2+^ influx is expected (apparent Ca^2+^-cooperativity of release (*m*) close to 1) when varying the number of open Ca^2+^-channels. Here, we found a near-linear dependence (operationally defined as *m* < 2) of glutamate release at single AZs on the whole-cell Ca^2+^-influx when depolarizing IHCs within the range of physiological receptor potentials (Supplementary Fig. 3a-c, *m*_*QCa*_ = 1.55, *n* = 29 synapses, N = 11 IHCs). Varying the voltage in this hyperpolarized range changes the open-channel number, and for Ca^2+^-nanodomain control, this is more relevant than the change in single-channel current as once the channel opens the ensuing Ca^2+^-signal tends to saturate the Ca^2+^ sensor of release^22^. In contrast, and consistent with the supralinear intrinsic Ca^2+^-dependence of exocytosis in IHCs^23,34^, reduction of the effective single-channel current by the rapid flicker-block of Ca^2+^-channels by Zn^2+^ showed *m* > 2 (Supplementary Fig. 3d-f, *m*_*QCa*_ = 2.56, *n* = 24 synapses, N = 10 IHCs). Taken together, imaging of glutamate release as a function of whole-cell Ca^2+^ influx corroborates the notion of a Ca^2+^-nanodomain-like control of release in mature IHCs^21–23,35^. However, the presynaptic heterogeneity of Ca^2+^-signaling in IHCs^11,20^ underscores the need of studying Ca^2+^ influx-release coupling at individual AZ.

Studying the Ca^2+^-dependence of exocytosis at individual AZs has remained difficult. Previous studies related whole-cell Ca^2+^-influx to either whole-cell exocytosis or to postsynaptic recordings of SV release. However, IHC synapses fundamentally vary in voltage dependence, number and clustering of their Ca^2+^-channels^11,19,20,28^. Here, we studied the operation of individual IHC synapses in the physiologically relevant voltage range during the 4^th^ postnatal week (P21-26). We combined ruptured-patch-clamp of IHCs with sequential dual-color imaging of first synaptic Ca^2+^-signals and then glutamate release (Fig. 2a). To image synaptic Ca^2+^-signals, we used the red-shifted, low-affinity chemical Ca^2+^-indicator Rhod-FF. We isolated Ca^2+^-signals at individual AZs using strong cytosolic buffering (10 mM EGTA in the patch pipette) and increased [Ca^2+^]_e_ (5 mM)^11,19,20^. We recorded glutamate release using ΔF-iGluSnFR in SGN boutons contacting the mid-section of IHC.

**Figure 2:**
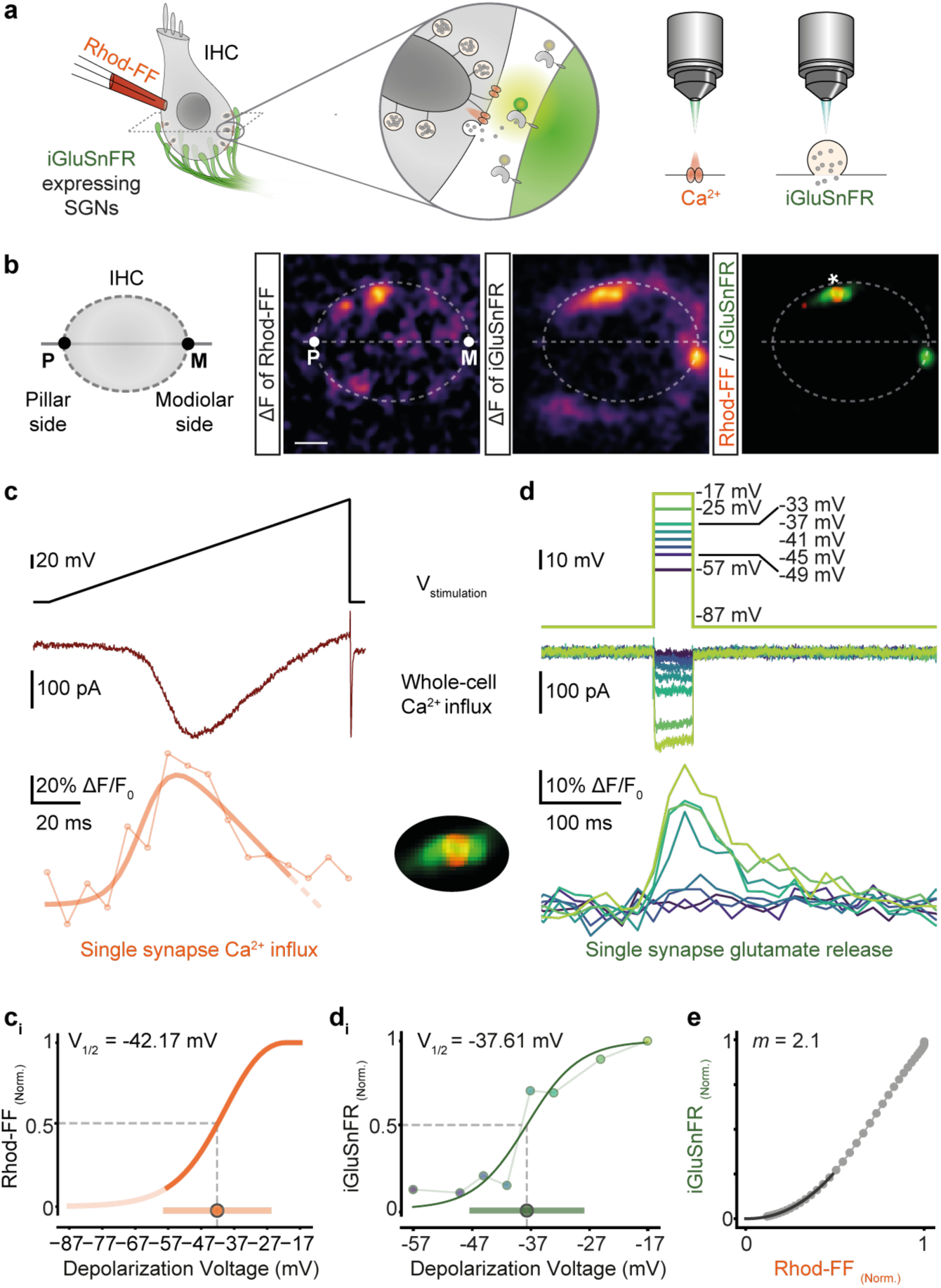
Sequential dual-color imaging of synaptic Ca^2+^-influx and glutamate release. **a.** IHCs were patch-clamped in ruptured-patch configuration with 800 μM Rhod-FF in the patch pipette, and simultaneously imaged for Ca^2+^-signals or glutamate release by spinning disc microscopy. **b.** Mean ΔF images of Rhod-FF and iGluSnFR signals in response to a voltage ramp and a 50-ms-long step depolarization, respectively. The synapse marked with * on the overlaid image is further analyzed in the following panels. (P: pillar side, M: modiolar side; Scale bar: 2 μm) **c** and **d.** Voltage command (top), corresponding whole-cell Ca^2+^-influx (middle) and the functional fluorescence responses (bottom) (bandstop filtered at 33.3 Hz) from Rhod-FF and iGluSnFR, respectively. A modified Boltzmann function (see Methods, R^2^ = 0.81) was fitted to a Rhod-FF fluorescence trace in response to a voltage ramp (c-c_i_). iGluSnFR-AUC was calculated per depolarization voltage and used for a Boltzmann fit (d_i_, R^2^ = 0.92). The voltage of half-maximal activation (V_1/2_) and the dynamic range (10-90%) of synaptic Ca^2+-^influx and glutamate release were calculated from the fits and depicted as circle and bar in c_i_ and d_i_, respectively. **e.** The obtained fits from a Ca^2+^ “hotspot” (c_i_) and from glutamate release (d_i_) were plotted against each other in a voltage range from −57 mV to −17 mV in 1 mV increments. A power function was fitted to the 25% of the maximum iGluSnFR-AUC (R^2^ = 0.99) to obtain the *m*-estimate. (ruptured patch-clamp, 10 mM intracellular EGTA, 5 mM [Ca^2+^]_e_) (See also Supplementary Fig. 4)

Next, we studied the voltage dependence of synaptic Ca^2+^-influx and glutamate release. To probe the voltage dependence of synaptic Ca^2+^-influx, we imaged Rhod-FF fluorescence (Fig. 2b, c) while applying a voltage ramp (−87 mV to +63 mV), a fast protocol inducing a minimum cellular Ca^2+^-load^11^. This allowed us to analyze the Ca^2+^-influx of the AZ corresponding to a given SGN bouton, by repeating the voltage ramps in five different planes (separated by 0.5 μm). The hotspots of Rhod-FF fluorescence elicited by depolarizations localized to the plasma membrane (Fig. 2b) and to the synaptic ribbon (Supplementary Fig. 4), indicating a cytosolic rise of [Ca^2+^] near the Ca^2+^-channel cluster of the AZ. Then, to probe the voltage dependence of glutamate release, we imaged iGluSnFR (on the central plane of Ca^2+^-imaging), while applying 50-ms-long step depolarizations (ranging from −57 to −17 mV) in a pseudo-randomized fashion (Fig. 2b, d). We used 50-ms-long depolarizations to elicit sufficient IHC release in the presence of 10 mM EGTA, which is expected to constrain the Ca^2+^-signal to the nanometer-proximity of the Ca^2+^-channels and to inhibit Ca^2+^-dependent SV replenishment^29^. Furthermore, the iGluSnFR-AUC seemed robust toward saturation up to at least 100 ms of stimulation (see Supplementary Fig. 1d-g). We analyzed the voltage-dependences of synaptic Ca^2+^-influx and glutamate release by fitting Boltzmann functions to ΔF/F_0_ of Rhod-FF (modified Boltzmann function, see Methods) and iGluSnFR-AUC. In the example shown, the voltages of half-maximal activation (V_1/2_) of Ca^2+^-influx (Fig. 2c_i_) and glutamate release (Fig. 2d_i_) were −42.2 and −37.6 mV, respectively. The resulting fit functions were then used to relate the glutamate release and the synaptic Ca^2+^-influx over the physiologically relevant voltage range. To estimate the apparent Ca^2+^-dependence of release for individual synapses, we fitted a power function on this relation (Fig. 2e). We restricted the fit until the 25% of maximum iGluSnFR-AUC for all synapses, in order to avoid saturation, e.g. due to RRP depletion and obtained an *m* estimate (*m =* 2.1 for the exemplary synapse). In conclusion, the sequential dual-color imaging approach allowed us to study Ca^2+^-dependence of release at individual AZs.

### IHC synapses vary in voltage dependence and apparent Ca^2+^-dependence of release

When systematically analyzing IHCs for the voltage dependence of whole-cell Ca^2+^-influx (Q_Ca_), synaptic Ca^2+^-influx (ΔF/F_0_ of Rhod-FF), and glutamate release (iGluSnFR-AUC), we observed fundamental heterogeneity of AZs within and across IHCs. The threshold for glutamate release was −48.27 ± 6.47 mV (SD, *n* = 55 synapses, N = 34 IHCs from 28 mice) and showed a broader distribution than that of the whole-cell Ca^2+^-influx (V_10_ = −47.16 ± 3.19 mV, SD, *p* < 0.0001, Levene’s test). As previously reported^11^, synaptic Ca^2+^ influx (V_1/2_ = −41.15 ± 5.70 mV, SD; *n* = 55 synapses, N = 34 IHCs from 28 mice, Fig. 3b) showed a broader and more negative V_1/2_ distribution than that of the whole-cell Ca^2+^-influx (V_1/2_ = −35.44 ± 2.83 mV, SD, Fig. 3a). The V_1/2_ distribution of synaptic glutamate release ranged from −45.25 mV to −29.86 mV and also showed a more negative average V_1/2_ (−37.49 ± 3.71 mV, SD; *n* = 55 synapses, *p =* 0.008, Fig. 3c) than the whole-cell Ca^2+^-influx. The V_1/2_ distributions differed significantly between glutamate release and synaptic Ca^2+^-influx (Supplementary Fig. 5a and *p* = 0.009, Levene’s test) as well as between synaptic and whole-cell Ca^2+^ influx (*p* = 0.001, Levene’s test). The V_1/2_ values of synaptic Ca^2+^-influx and glutamate release correlated (Pearson’s *r* = 0.43, *p* = 0.0008, Fig. 3g). This correlation indicates that the differences in the voltage dependence of Ca^2+^ influx are propagated to release, generating heterogeneous output among the IHC AZs at a given receptor potential.

**Figure 3.**
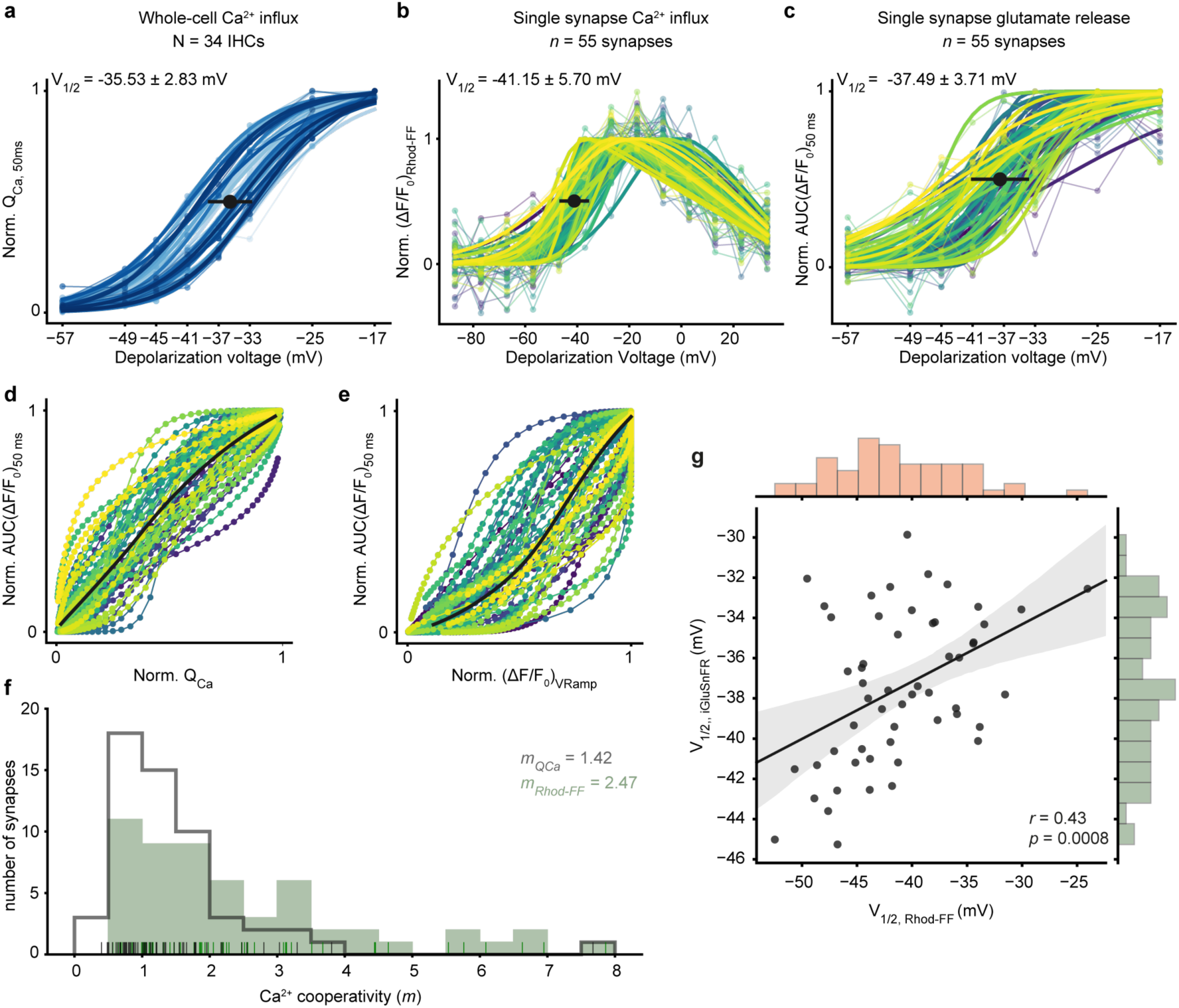
IHC synapses vary in voltage dependence and apparent Ca^2+^-dependence of release. **a.** Normalized whole-cell Q_Ca_, calculated in response to 50-ms-long step depolarizations, is plotted as a function of depolarization voltage. A Boltzman function was fitted to estimate the V_1/2_. Individual IHCs are color coded in shades of blue (mean ± SD, N = 34 IHCs from 28 mice). **b.** A voltage ramp was applied to obtain ΔF of Rhod-FF as a proxy synaptic Ca^2+^ influx. The V_1/2_ of synaptic Ca^2+^-influx was calculated from a modified Boltzman function (see Methods) fitted to ΔF/F_0_. (mean ± SD, *n* = 55 synapses; individual synapses are color coded). **c.** Normalized iGluSnFR-AUC, in response to 50-ms-long step depolarizations, same as A. **d.** The relation of whole-cell Ca^2+^-influx (a) and the synaptic glutamate release (c). **e.** The relation of synaptic Ca^2+^-influx (b) and glutamate release (c). The bold lines show the means. (See also Supplementary Fig. 6 for individual plots) **f.** The histogram shows the Ca^2+^-cooperativities (*m*) obtained by individual power function fitting until 25% of normalized iGluSnFR response from (d; gray) and from (e; green). The mean *m* was found to be 1.42 and 2.47, respectively. The rug plot shows the individual data points. **g.** The V_1/2_ of synaptic Ca^2+^-influx and glutamate release are correlated (Pearson’s *r* = 0.43, *p* = 0.0008). The marginal histograms show the distribution of each axis. Linear regression analysis (solid lines) and the associated 95% confidence intervals (shaded area). (See also Supplementary Figs. 3-5)

Next, we estimated the apparent Ca^2+^-dependence of release by relating the iGluSnFR-AUC to the whole-cell Ca^2+^-influx (Fig. 3d) and to the synaptic Ca^2+^-influx (Fig. 3e) for the above experiments that primarily varied the open-channel number (see also Fig. 1c-d and Supplementary Fig. 3). The apparent Ca^2+^-dependence of glutamate release at the single-synapse level (Fig. 3f) was higher on average (*m*_*Rhod-FF*_ = 2.47) than when relating glutamate release to the whole-cell Ca^2+^-influx (*m*_*QCa*_ = 1.42). The latter was lower on average than the *m* estimate obtained in the perforated-patch configuration on P15-19 IHCs at 1.3 mM [Ca^2+^]_e_ using a similar stimulus protocol (see Fig. 1c-d; Supplementary Fig. 3a-c, *p =* 0.001, Mann-Whitney-U-Test). This difference is compatible with the hypothesis that strong Ca^2+^ buffering by 10 mM EGTA favors the operation of release sites under Ca^2+^-nanodomain control. Since the data of Fig. S3 were acquired at an earlier developmental stage (P15-19), the developmental tightening of the Ca^2+^ channel-exocytosis coupling^23^ might also have contributed to the lower *m* in Fig. 3d (P21-26). When operationally defining *m* < 2 (“near-linear”) as indicative of Ca^2+^-nanodomain-like control of release, we found approximately half of the synapses (29 out of 55 synapses) to operate in this scenario. The other half showed a broad spread of *m* values reaching up to 8, compatible with Ca^2+^-microdomain-like control of release despite the presence of 10 mM EGTA. Taken together, single-synapse imaging of Ca^2+^-influx and glutamate release revealed a heterogeneity of the apparent Ca^2+^-dependence of release that is likely due to differences of the coupling of Ca^2+^-channels to release-sites among the IHC AZs. This heterogeneous coupling of Ca^2+^-channels to release sites is likely one of the underlying mechanisms contributing to the heterogenous output of IHC AZs.

### Pillar synapses operate at more negative potentials than modiolar synapses

SGNs exhibit a spatial preference in their IHC innervation pattern in the cat: high SR-low threshold fibers innervate on the pillar side of the IHC, and low SR-high threshold fibers contact the modiolar side of the IHC^6^. Assuming analogy for mouse IHCs, we probed how presynaptic heterogeneity might contribute to the functional diversity of SGNs by analyzing the synaptic Ca^2+^-influx and glutamate release as a function of position along the pillar-modiolar axis. Fig. 4a compares the simultaneous iGluSnFR responses of two exemplary synapses of one IHC: one positioned at the pillar and the other one at the modiolar side. In this example, the pillar synapse had a lower threshold, i.e. it already became active at −49 mV, while the modiolar synapse only started to respond at −41 mV. The pillar synapse also showed a more negative V_1/2_ and a wider dynamic range compared to the modiolar synapses in the given cell (Fig. 4a_i_). As this observation was made in the same section of an IHC and was representative for the population (Fig. 4b-e), we consider potential technical reasons unlikely. On average, pillar synapses exhibited a more negative activation threshold as well as V_1/2_ of both Ca^2+^-influx and glutamate release (Fig. 4b-d). Moreover, pillar synapses showed a wider dynamic range of release (Fig. 4e). Nonetheless, there was substantial variability in particular among the modiolar synapses. This is obvious from the example shown in Fig. 4a_i_ as well as at the population level (Fig. 4b-e). We also performed linear regression analysis on the relation of pillar-modiolar position and V_1/2_ of synaptic Ca^2+^-influx as well as glutamate release (Fig. 4b-e). This analysis confirmed a pillar-modiolar gradient of V_1/2_ of synaptic Ca^2+^-influx (*r* = 0.48; *p* = 0.0001) that we previously reported for an earlier postnatal stage (P14-20)^11^. Furthermore, it showed a pillar-modiolar gradient of the voltage threshold (*r* = 0.47; *p* = 0.0002), V_1/2_ (*r* = 0.32; *p* = 0.013) and dynamic range (*r* = −0.38; *p* = 0.003) of release.

**Figure 4.**
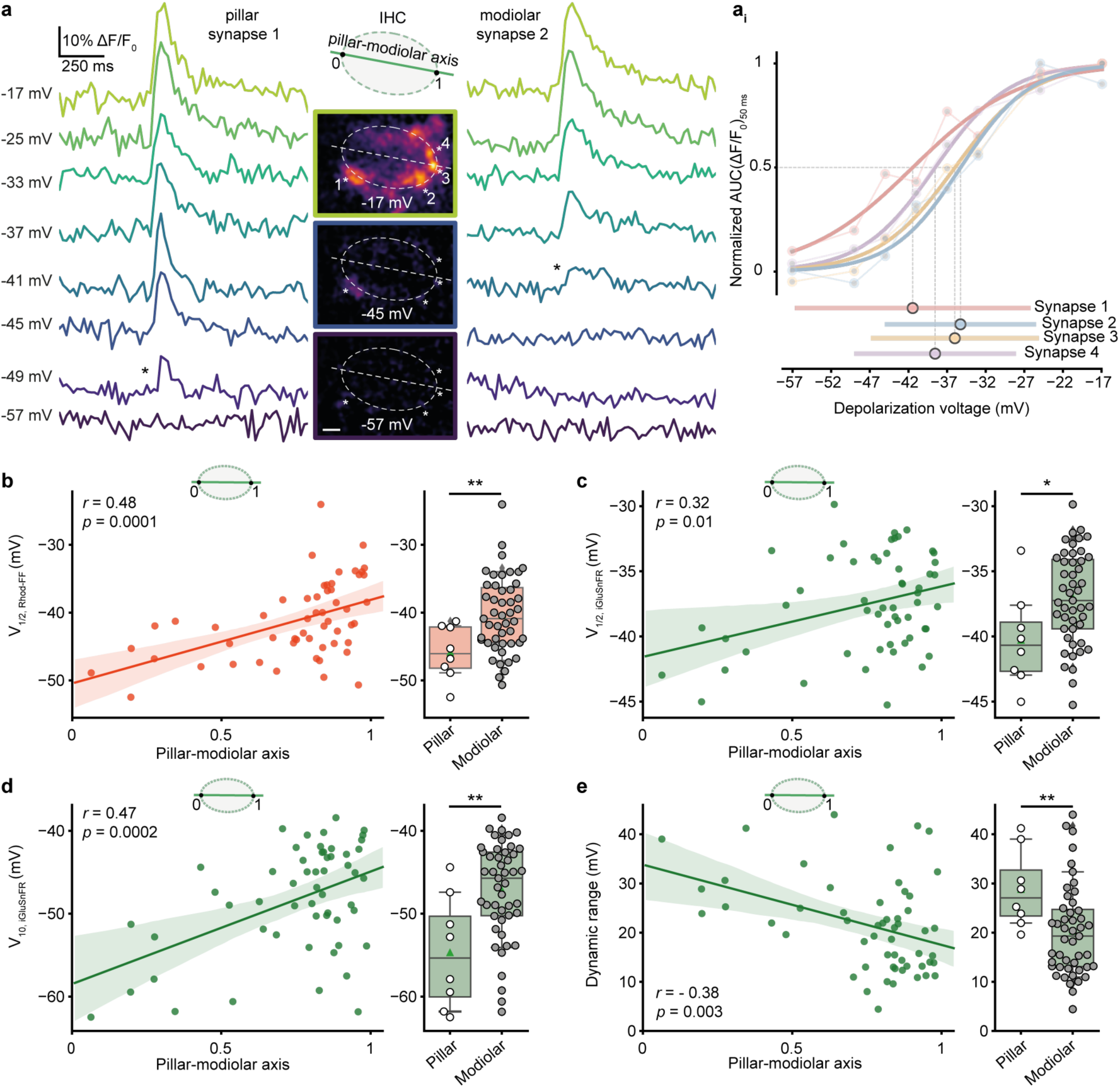
Pillar synapses are active at more negative potentials than modiolar synapses. **a.** This single cell example shows release dynamics of two synapses innervating the same IHC from either pillar or modiolar side. iGluSnFR signals (bandstop filtered at 33.3 Hz) in response to 50-ms-long step depolarizations to the given depolarization voltages are depicted. The middle panel shows ΔF images of iGluSnFR, recorded from the mid-section of the IHC. * depict the first detected response in the given synapse. Note that pillar synapse 1 is already active at −49 mV, while modiolar synapse 2 starts responding only at −41 mV. (Scale bar 2 μm) a_i_ shows the normalized iGluSnFR-AUC as a function of depolarization voltage. Dynamic range and V_1/2_ of the synapses are depicted in the lower panel. **b-e.** Left panel shows the linear regression analysis (solid lines) of V_1/2_ of synaptic Ca^2+^-influx (b), glutamate release (c), threshold (d) and dynamic range of release (e) as a function of position along the pillar-modiolar axis. Shaded areas depict the associated 95% confidence intervals. Right panel shows box and whisker plots of these properties of synapses grouped into pillar and modiolar halves of the IHCs. * *p* ≤ 0.05, ** *p* ≤ 0.01, *** *p* ≤ 0.001

In conclusion, pillar and modiolar AZs differed in their voltage dependence and dynamic range of glutamate release. These differences between pillar and modiolar synapses support the hypothesis that diversity of the spontaneous and sound-evoked SGN firing is, at least in part, routed in the heterogenic biophysical properties of presynaptic Ca^2+^-channels and transmitter release. Activation at more negative potentials of pillar synapses agrees with the high SR and low sound threshold of the corresponding SGNs. However, the wider dynamic range of glutamate release at pillar AZs is more difficult to reconcile with the narrower dynamic range of sound encoding of high SR-low threshold SGNs^5^.

### Clustering of synaptic properties indicates three synapse subtypes likely distinguished by their implementation of Ca^2+^ influx-release coupling

By the sequential dual-color imaging of synaptic Ca^2+^-influx and release, we quantified each synapse with 15 parameters. Correlations among several of these parameters can be intuitively explained, such as the voltage-dependence of a synapse, which is primarily rooted in that of Ca^2+^-channel activation (Fig. 5). Moreover, a supralinear coupling of release to Ca^2+^-influx is expected to compress the dynamic range of release. While the comparative analysis of pillar and modiolar synapses (see Fig. 4) aims to elucidate synaptic correlates of the functional SGN properties, it has both value and limits. For an unbiased and in-depth analysis, we applied K-means clustering (K = 3) on the 11-th dimensional space of single-synapse properties (excluding positional and whole-cell information) and obtained 3 synapse clusters (putative subtypes). To visualize the clusters in two or three dimensions, we performed principal component analysis (PCA) and used the first three principal components that explained 79% of the variance (Fig. 6a-d).

**Figure 5:**
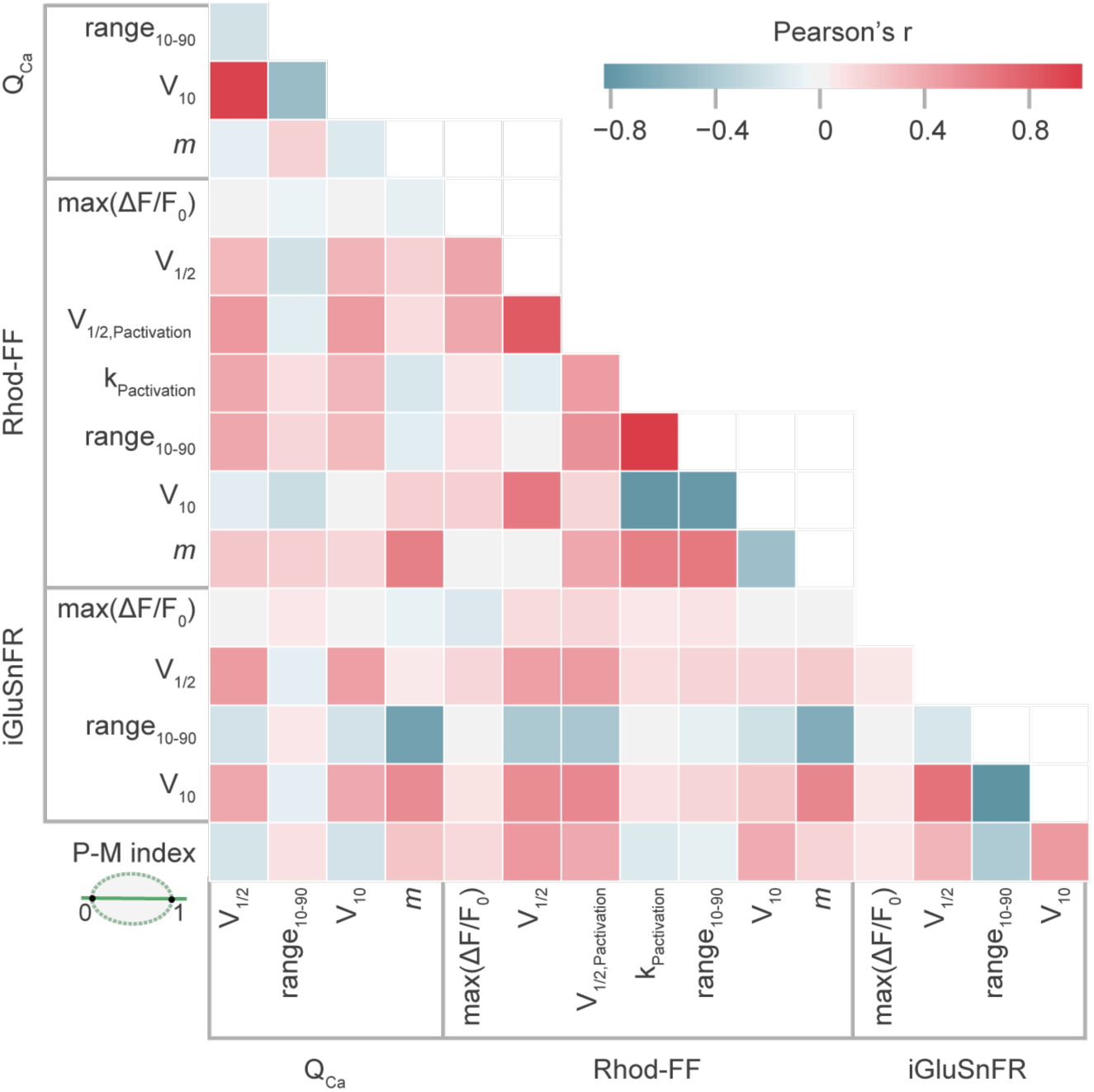
Correlation map of synaptic properties. This correlation matrix shows the Pearson correlation coefficients (Pearson’s *r*) between various properties assigned to individual synapses. The degree of correlation is color-coded: light (weak) to dark (strong). Positive correlations are depicted in red and negative ones in blue.

**Figure 6.**
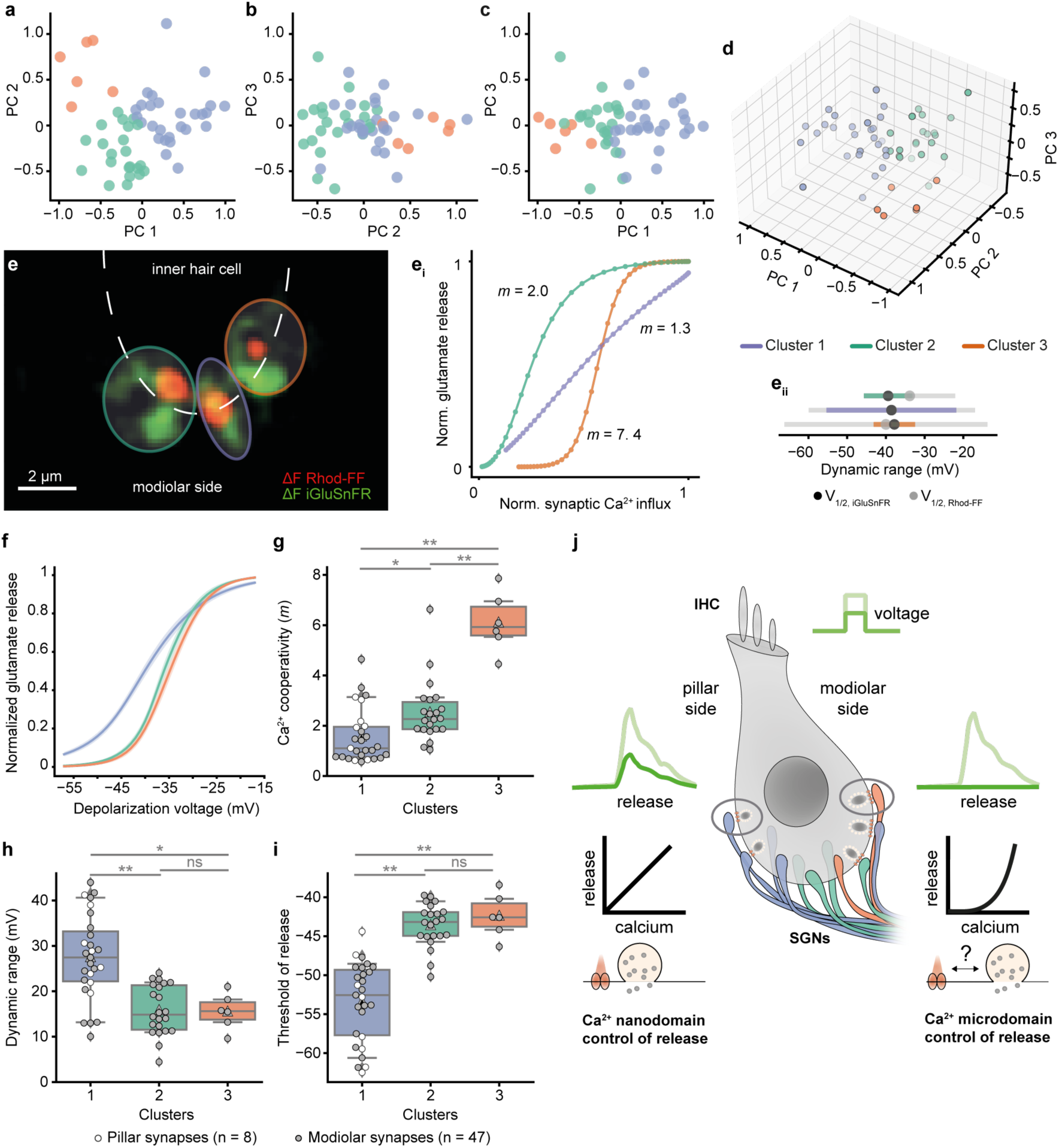
Three putative synapse subtypes, co-existing in individual IHCs, differ in their synaptic transfer functions and Ca^2+^-dependencies. 11 synaptic properties (except for the positional information; see Fig. 5) of 55 synapses (N = 34 IHCs from 28 mice) were used for the PCA and K-means clustering. **a, b** and **c** show the 2D plots of the first 3 principal components (PCs), labeled based on the clusters obtained by K-means clustering algorithm. **d.** 3D plot showing the first 3 PCs. **e.** Single IHC exhibits different modes of Ca^2+^-control of release. The overlaid ΔF image of Rhod-FF (red) and iGluSnFR (green) shows synaptic Ca^2+^-influx and glutamate release at three neighboring modiolar synapses (individual synapses are color coded based on their clusters). **e**_**i**_. The relation between Ca^2+^-influx and glutamate release of given synapses in the voltage range of −57 to −17 mV plotted with 1 mV increments. A power function was fitted until the 25% of normalized iGluSnFR AUC(ΔF/F_0_). Synapses showed different Ca^2+^-dependencies (*m* = 2.0, 1.3, 7.4; green, purple, orange). **e**_**ii**_. Dynamic ranges of corresponding Ca^2+^-influx (gray) and glutamate release (color-coded) with V_1/2_ depicted. Note that the synapses from cluster 1 (purple), employing Ca^2+^-nanodomain-like control of release (*m* = 1.3), exhibit wider dynamic range than the other synapses. **f.** Mean glutamate release (iGluSnFR-AUC) as a function of depolarization voltage in the three identified clusters (mean ± SEM). The Ca^2+^-cooperativity (*m*) (**g**), dynamic range (**h**), and glutamate release threshold (V_10_) (**i**) show differences for the three clusters. Cluster 1 is composed of linear synapses with wider dynamic range and lower threshold compared to the other clusters. Pillar synapses are depicted as white-filled circles, and modiolar ones are depicted as gray-filled circles. The clusters were compared by one-way ANOVA test, followed by a posthoc Tukey’s test. * *p* ≤ 0.05, ** *p* ≤ 0.01, *** *p* ≤ 0.001 **j.** The proposed model for sound intensity encoding in an IHC. Differences in the presynaptic control of release, in terms of Ca^2+^-signaling and Ca^2+^ channel-exocytosis coupling, enable IHC to diversify the transfer functions of individual synapses for the same receptor potentials.

Next, we checked the response properties of these putative synapse subtypes. The first subtype of synapses showed a low *m* (1.56 ± 1.07, 27 synapses) indicating a Ca^2+^-nanodomain-like control of release as well as a wide dynamic range (27.52 ± 9.37 mV) and a hyperpolarized activation threshold (−53 ± 5.07 mV) of release. In contrast, the third synapse subtype showed a high *m* (6.10 ± 1.18, 6 synapses) with smaller dynamic range (15.53 ± 3.95 mV) and a more depolarized threshold (−42 ± 2.8 mV) (Fig. 6f-i). Synapses from the first cluster are predicted to be active already around the IHC receptor potential (−55 mV, Fig. 6i)^32^. The properties of the second synapse subtype fell in between those of subtypes 1 and 3. Finally, we evaluated positions of the three synapse subtypes in an effort to match them to the concept of a SGN-subtype specific innervation pattern along the pillar-modiolar axis. Pillar synapses were exclusively of subtype 1, while modiolar synapses were distributed across all subtypes. This is exemplified by three neighboring modiolar synapses of an IHC (Fig. 6e): each synapse belongs to a separate cluster, employing different apparent Ca^2+^-dependencies (see Supplementary Fig. 6 for the data of all individual IHCs). Hence, different topographies of Ca^2+^-channels and SV release sites apparently co-exist even in compact presynaptic cells. How the IHC manages to vary the AZ organization and, thereby, Ca^2+^ influx-exocytosis coupling remains to be elucidated. Our data indicate some extent of pillar-modiolar segregation of synaptic properties, while substantial heterogeneity is found among the AZ at all positions of the synaptic IHC pole. Taken together, we propose a model in which IHCs fractionate the coding of sound intensity information, contained in the receptor potential, via heterogenous synaptic input-output functions that are sourced by differences in voltage dependence and release-site coupling to Ca^2+^-channels (Fig. 6j).

## Discussion

The auditory system encodes sound pressures over a range of six orders of magnitude. The sensory mechanisms contributing to this fascinating behavior include active cochlear micromechanics, adaptation at various stages, as well as diversity of SGNs and their synapses with IHCs. At each tonotopic position in the cochlea, SGNs differ in their spontaneous and evoked firing, effectively tiling the range of audible sound pressures with their responses^1–5^. Differences in the transcriptomic profiles of SGNs have led to their categorization into molecular subtypes^16–18^. Moreover, heterogeneity in the pre- and postsynaptic properties of afferent IHC-SGN synapses as well as of efferent SGN innervation have been proposed to explain the fractionation of the entire range of audible sound pressures, detected by the same IHC, into different neural representations^10,11,19,20,36^. Here, we studied the transfer at individual IHC-SGN synapses, by imaging of synaptic Ca^2+^-signals and glutamate release during IHC patch-clamp recordings. On average, glutamate release had a voltage threshold near the physiological resting potential (Fig. 1 and 4) and showed a near-linear dependence on the whole-cell Ca^2+^-influx when primarily varying the open-channel number (Supplementary Fig. 3). We then employed sequential dual-color imaging of Ca^2+^-signals and glutamate release of single AZs (Fig. 2). This way we revealed the heterogeneity of synaptic transfer functions, rooted in differences in voltage-dependence of the Ca^2+^-influx and in its coupling to exocytosis (Fig. 3). We found a pillar-modiolar gradient of AZ transfer functions: consistent with preferred innervation of high-SR SGNs on the pillar side, pillar AZs operated at more negative potentials than the modiolar ones (Fig. 4). By K-means clustering of AZs according to the single-synaptic parameters, we obtained three putative synapse subtypes, differing in the voltage-dependence, apparent Ca^2+^-dependence, and dynamics of release (Fig. 6).

### Heterogeneity of Ca^2+^ channel-release coupling

Studying Ca^2+^-signals and glutamate release at individual AZs revealed that the coupling of Ca^2+^-influx to exocytosis varies among the AZs, unlike previously assumed based on average AZ behaviour^21–23,35^. We show that it ranges from Ca^2+^-nanodomain to Ca^2+^-microdomain control of exocytosis and differs even among neighboring IHC AZs. Approximately one half of the AZs showed Ca^2+^-nanodomain-like control of exocytosis (operationally defined as *m* < 2) during primary manipulation of the open-channel number. Hence, different from the summation model^24^, Ca^2+^-nanodomain-like control of exocytosis occurs at individual IHC AZ. However, we found the other half of the AZs with *m >2* suggesting Ca^2+^-microdomain control of exocytosis to coexist. We assume all AZs share the same intrinsic Ca^2+^-cooperativity of 4-5 that was established for IHC exocytosis by C_m_ recordings^21,34^. This likely reflects the cooperative binding of 4-5 Ca^2+^ ions to the putative Ca^2+^-sensor of IHC release, otoferlin^37^. In the 4^th^ postnatal week, we consider IHCs to have reached maturity but further changes might occur^38^. However, the heterogeneity was preserved in all recorded ages (P21-26, data not shown). Here, we studied synapses of apicocochlear IHCs and, given the previously described tonotopic differences in Ca^2+^ channel-release coupling on the whole-cell level^39^, single-synapse analysis in basocochlear IHCs remains an important task for the future. Taken together, we propose a model by which IHCs vary the topographies of Ca^2+^-channels and SV release sites at the AZs likely to diversify synaptic transmission beyond the heterogeneity of AZ size and Ca^2+^-signaling.

Non-exclusive candidate mechanisms for position-dependent AZ diversification in IHCs include i) developmental competition of synapses, ii) transsynaptic signaling from SGNs, and iii) planar polarity signaling. One interesting idea is that, during development, pioneer SGN axons^40^ making the first synaptic contact at the modiolar side of IHCs attract large amounts of presynaptic resources. In addition to such potential transsynaptic cueing of AZ properties, IHCs might employ their planar polarity signaling to instruct modiolar-pillar gradients of AZ size^41^. How such signaling differentially shapes AZ properties remains an exciting topic for future studies. Clearly, polarized trafficking of components of the Ca^2+^-channel complex or other AZ proteins could contribute. This might, for instance, involve the adapter and PDZ-domain protein Gipc3, defective in human deafness^42^, required for the modiolar-pillar gradient of maximal synaptic Ca^2+^-influx^11^. Intriguingly, the spatial distribution of the number of Ca^2+^-channels and their voltage dependence might be regulated by different mechanisms^41^. While the number of Ca^2+^-channels scales with AZ/ribbon size^11,19,20^ and shows a modiolar-pillar gradient, the topography relative to SV release sites and biophysical properties of the Ca^2+^-channels seem to follow an opposing gradient. For instance, the voltage-dependence of synaptic Ca^2+^-influx^11,41^ and, consecutively, glutamate release (Fig. 4) shows a pillar-modiolar gradient, with activation at more hyperpolarized potential for the pillar AZs that show smaller Ca^2+^-channel clusters. Likewise, *m* of exocytosis was typically lower at the pillar side, indicating that Ca^2+^-nanodomain control of release prevails with the lower number of Ca^2+^-channels, while an increased channel number favors domain overlap as also predicted by modeling^23^. Future studies will need to probe for differences in the abundance of potential molecular linkers^43–46^ or spacers of Ca^2+^-channels and release sites such as RIM and RIM-BP among the IHC AZs and to explore actual Ca^2+^ channels-release site topography of IHC AZs e.g. by electron microscopy (ref.^47^).

As for size and Ca^2+^-channel number of the AZ, the position-dependence of biophysical properties and topography of Ca^2+^-channels might be shaped during postnatal development. For instance, AZs of immature IHCs, on average, employ a Ca^2+^-microdomain-like control of exocytosis. Maturation tightens the coupling of Ca^2+^-influx and exocytosis at least when considering the collective behavior of all IHC AZs^23^, coinciding with the appearance of high-SR fibers^48^. This study indicates that a subset of mature IHC synapses employs Ca^2+^-microdomain control, representing an additional mechanism employed by the IHC to diversify synaptic transmission and endowing this subset with further potential of presynaptic plasticity^49^.

Potential pitfalls of our imaging approach to single IHC AZ function include: i) glutamate spill-over from neighboring synapses, ii) saturation of iGluSnFR and other non-linear effects resulting from the glutamate binding to endogenous receptors in the postsynapse, and iii) contamination of synaptic Ca^2+^-signals by mitochondrial-Ca^2+^ changes, as some rhodamine-based dyes partition into mitochondria^50^. We aimed to minimize synaptic crosstalk by choosing well-separated boutons and region of interests (ROIs) with a small radius (<1 µm)^51^. While we cannot rule out a contribution of mitochondrial Rhod-FF, the use of 10 mM EGTA would make mitochondrial-Ca^2+^ uptake unlikely. This is supported with the strict co-localization of the Rhod-FF signals with the synaptic ribbon (Supplementary Fig. 4).

### Relating presynaptic heterogeneity to functional SGN diversity

The functional subtypes of SGNs are said to spatially segregate their IHC innervation: high-SR SGNs preferentially innervate the pillar side of the IHC, while low-SR SGNs preferentially innervate the modiolar side of the IHC in the cat cochlea^6^. Here, we demonstrate by iGluSnFR imaging in the apical organ of Corti of mice that glutamate release from pillar synapses operates at more hyperpolarized potentials than the modiolar ones. This offers an exciting presynaptic hypothesis for the functional SGN diversity: resting potential or weak receptor potentials will primarily recruit pillar synapses, which can readily explain the high SR and low sound threshold of SGNs innervating the pillar IHC side. While this hypothesis was previously phrased based on the heterogeneous voltage-dependence of Ca^2+^-influx^11^, whether and how this translates into heterogeneity of glutamate release remained unclear. Indeed, unlike we had previously assumed^11^, the present study revealed differences in the Ca^2+^ channel-release coupling among the IHC AZs. Heterogeneity of Ca^2+^ channel-release coupling, too, could contribute to diversifying SGN function. First, Ca^2+^-nanodomain control of exocytosis would increase the SR, as stochastic opening of Ca^2+^-channels could trigger release in such a tight coupling scenario^52,53^. Second, Ca^2+^-nanodomain control of exocytosis likely also promotes low voltage threshold for the same reason and, indeed, we found a correlation between the threshold of release and *m* (Fig. 5). Third, Ca^2+^-nanodomain control of exocytosis widens the dynamic range of release. Indeed, we found a negative correlation between dynamic range and *m* (Fig. 5). One of the most puzzling questions is, how the above hypothesis of pillar AZs with more hyperpolarized operation driving high-SR SGNs that show a smaller dynamic range of sound encoding can be reconciled with the wider dynamic range of glutamate release found for pillar AZs. One possible explanation is high spontaneous release at the *in vivo* IHC resting potential lowers the standing RRP available for sound evoked release and hence causes a narrower dynamic range of release (reviewed in ref.^54^). Indeed, disruption of PDZ protein Gipc3 resulted in enhanced SR and narrower dynamic range^11^. Furthermore, as rate-level functions are usually assessed in response to sound stimuli lasting more than 50 ms, there is likely a contribution of RRP replenishment^22^. We cannot exclude possible contribution of postsynaptic properties, efferent modulation, dynamic range adaptation^55^, and non-linearity imposed by the basilar membrane^4^ on the dynamic range of SGN firing. Lastly, differences in Ca^2+^ channel-exocytosis coupling could affect the short-term plasticity (ref.^56^), note the higher adaptation strength of high-SR SGNs *in vivo* (ref.^57^).

In conclusion, we propose a model where differences among the IHC AZs in the presynaptic control of release, in terms of presynaptic Ca^2+^-signaling and Ca^2+^ channel-exocytosis coupling, enable the IHC to diversify the synaptic signaling to SGNs for the same receptor potential.

## Acknowledgements

We thank Dres. Christian Vogl and Vladan Rankovic for their contributions in the initial phase of the project, N. Dietrich, S. Gerke, and C. Senger-Freitag for expert technical assistance. We thank Dr. G. Ramos-Traslosheros for his initial assistance with the analysis. We thank Dr. Thomas Euler for providing us with a sample of AAV9 *hSyn*.iGluSnFR. We thank Dres. J. Neef, L.M. Jaime Tobón, E. Neher and S.O. Rizzoli for helpful comments on the manuscript. We thank Dr. Vladimir Belov for providing Abberior Star 488-conjugated peptide. Ö.D.Ö. has been a doctoral student of the Ph.D program “Neurosciences” – International Max Planck Research School and the Göttingen Graduate Center for Neurosciences, Biophysics, and Molecular Biosciences (GGNB) (DFG grant GSC 226) to the Georg August University Göttingen. This work was supported by the Deutsche Forschungsgemeinschaft (DFG, German Research Foundation) under Germany’s Excellence Strategy – EXC 2067/1-390729940 to T.M. as well as by the DFG’s Leibniz Program to T.M.

## Author Contributions

T.M and Ö.D.Ö designed the study. Ö.D.Ö performed the experiments and the analysis. T.M and Ö.D.Ö prepared the manuscript.

## Methods

### Animals and postnatal injections

All experiments were done in compliance with national animal care guidelines and were approved by the University of Göttingen Board for Animal Welfare and the Animal Welfare Office of the State of Lower Saxony (permit number: 17-2394). Postnatal AAV-injections were made into scala tympani of the right ear through the round window^58,59^. P5-7 WT C57Bl/6 mice were used for the injection of AAV9 virus under human synapsin promoter (pAAV9.*hSyn*.iGluSnFR, commercially available from Addgene, USA) to drive transgenic expression of iGluSnFR in SGNs. In brief, under general and local anesthesia (isoflurane and xylocaine, respectively), the right ear was accessed through a dorsal incision. Once the round window membrane was located, a quartz capillary pipette was used to gently puncture it and inject ∼1-1.5 μl of pAAV9.*hSyn*.iGluSnFR (titer ≥ 1 × 10^13^ vg/ml). After the injection, the wound was sutured and buprenorphine (0.1 mg/kg) was applied as a painkiller. The recovery of the animals was monitored on a daily basis. All animals were kept in a 12-hour light/dark cycle, with access to food and water *ad libitum* and together with the mother until the end of the weaning period (∼P21). Injected WT mice were used for experiments either one week (P15-19) or two weeks (P21-26) after the injection.

### Auditory Brainstem Recordings

Recordings of auditory brainstem responses (ABR) were performed on P29 mice as previously described^60^. Briefly, mice were anesthetized with a combination of ketamine (125 mg/kg) and xylazine (2.5 mg/ kg) intraperitoneally. The core temperature was maintained constant at 37°C using a heat blanket (Hugo Sachs Elektronik–Harvard Apparatus). The TDT II system run by BioSig software (Tucker Davis Technologies) or by MATLAB (MathWorks) was used for stimulus generation, presentation, and data acquisition. Tone bursts (6/12/24 kHz, 10 ms plateau, 1 ms cos^2^ rise/fall) or clicks of 0.03 ms were presented at 40 Hz (tone bursts) or 20 Hz (clicks) in the free field ipsilaterally using a JBL 2402 speaker.

### Immunohistochemistry and confocal microscopy

Acutely dissected apical turns of organs of Corti were fixed in formaldehyde (4% in phosphate buffered saline (PBS), 1h on ice). After 3×5 min PBS washing steps, the samples were blocked with a goat serum dilution buffer (16% normal goat serum, 450 mM NaCl, 0.3% Triton X-100, 20 mM phosphate buffer, pH 7.4) for 1 hour at room temperature in a wet chamber. The blocking was followed by an overnight incubation with the primary antibodies at 4°C. After 3×5 min PBS washing steps, the samples were incubated with the secondary antibodies for 1 hour at room temperature. Following the final 4×5 min PBS washing steps, the samples were mounted in mounting medium (Mowiol 4-88, Sigma). The primary antibodies used were the following: mouse anti-CtBP2 (1:200, BD Biosciences, 612044) –to detect synaptic ribbons–, chicken anti-GFP (1:200, Abcam, 13970) –to detect iGluSnFR–, and guinea pig anti-parvalbumin (1:200, Synaptic Systems, 195004) –to detect SGNs, OHCs, and IHCs. Secondary goat antibodies were used with 1:200 dilution: Alexa Fluor 488 conjugated anti-chicken (Dianova, 703-45-155), Alexa Fluor 633 conjugated anti-mouse (Invitrogen, A31571), Alexa Fluor 488 anti-chicken (Invitrogen, A11039) and Alexa Fluor 568 anti-guinea pig (Invitrogen, A11075). Images were acquired using an Abberior Instruments Expert Line STED microscope, with excitation lasers at 488, 561, and 640 nm using a 1.4 NA 100x or 20x oil immersion objective, in confocal mode. Images were adjusted for brightness and contrast using Image J for illustration purposes.

### Patch-clamp recordings

The apical 2/3 turn of organs of Corti were acutely dissected from P15 to P26 animals in HEPES Hanks’ solution containing (in mM): 5.36 KCl, 141.7 NaCl, 10 HEPES, 0.5 MgSO_4_, 1 MgCl2, 5.6 D-glucose, and 3.4 L-glutamine (pH 7.2, ∼300 mOsm/l). The IHC basolateral membranes were exposed by cleaning of nearby cells with a suction pipette by approaching from either pillar side or modiolar side. All experiments were conducted at room temperature (20-25°C). Patch pipettes were made from GB150-8P or GB150F-8P borosilicate glass capillaries (Science Products, Hofheim, Germany) for perforated and ruptured patch-clamp recordings, respectively. To decrease the capacitive noise, pipettes were coated with Sylgard and their tips were polished with a custom-made microforge. All patch-clamp recordings were done simultaneously with fluorescent imaging of iGluSnFR or of iGluSnFR and Ca^2+^.

#### Perforated-patch recordings

Perforated-patch clamp was performed as described previously^29^. For Ca^2+^ current and membrane capacitance (C_m_) measurements, the extracellular solution contained the following (in mM): 110 NaCl, 35 TEA-Cl, 2.8 KCl, 1 MgCl_2_, 1 CsCl, 10 HEPES, 1.3 CaCl_2_, and 11.1 D-glucose (pH 7.2, ∼305 mOsm/l) and was introduced into the recording chamber via a perfusion system. The pipette solution contained (in mM): 130 Cs-gluconate, 10 TEA-Cl, 10 4-AP, 10 HEPES, 1 MgCl2, as well as 300 mg/ml amphotericin B (pH 7.17, ∼ 290 mOsm/l). The intracellular solution also contained the TAMRA-conjugated dimeric CtBP2/RIBEYE-binding dimer peptide (10 μM, Biosynthan, Germany)^23,61^. To label the synaptic ribbons, the peptide was introduced to the cell by rupturing the membrane patch towards the end of recordings. The intracellular exposure to amphotericin (used to perforate the membrane for electrical access) did not result in a noticeable increase of IHC conductance and compromise IHC health. All the measurements were done via EPC-10 amplifiers controlled by Patchmaster software (HEKA Elektronik, Germany). The holding potential was −87 for all the recordings. All voltages were corrected for liquid junction potential offline (17 mV). Currents were leak-corrected using a p/10 protocol. Recordings were used only if the leak current was lower than 30 pA and the series resistance (Rs) was lower than 30 mOhm. The Lindau-Neher technique was used to measure the C_m_ changes^62^. Exocytosis was quantified from C_m_ changes as described previously^29,63^. IHCs were stimulated by step depolarizations of different durations (2 to 100 ms, applied in a pseudo-randomized manner) to −23 mV at intervals of 60s-100s (Supplementary Fig. 1d-g). To probe voltage-dependence of release, IHCs were step-depolarized for 10 ms to potentials ranging from −62 mV to −22 mV in 5 mV increments in a pseudo-randomized order (Fig. 1; Supplementary Fig. 2). For the Zn^2+^ perfusion experiments, 1 mM Zn^2+^ was slowly perfused in and out of the recording chamber, while IHCs were step depolarized for 10 ms to −23 mV simultaneously up to 20 times (Supplementary Fig. 1a-b).

#### Ruptured-patch recordings

Ruptured-patch experiments were performed in extracellular solution containing (in mM): 2.8 KCl, 102 NaCl, 10 HEPES, 1 CsCl2, 1 MgCl2, 35 TEA-Cl, 2 mg/ml D-Glucose and 5 CaCl_2_ (pH 7.2, 300 mOsm). The patch pipette solution contained (in mM): 111 L-glutamate, 1 MgCl_2_, 1 CaCl_2_, 10 EGTA, 13 TEA-Cl, 20 HEPES, 4 Mg-ATP, 0.3 Na-GTP and 1 L-Glutathione (pH 7.3, ∼290 mOsm). For fluorescent imaging, 800 μM of the low affinity chemical Ca^2+^ indicator Rhod-FF tripotassium salt (Kd:19 μM, AAT Bioquest, USA) was added to the intracellular solution. The recordings were discarded when the leak current exceeded −50 pA at −87 mV or R_S_ was greater than 15 MΩ within 4 min after break-in. For Ca^2+^ imaging experiments, a voltage ramp (from −87 to +63 mV during 150 ms; 1 mV/ms) was applied to evoke Ca^2+^ influx (Fig. 2,3).

For all IV recordings, the IHCs were step depolarized for 20 ms from −87 mV to +63 mV in 5 mV increments. The IV recordings were used to assess the fitness of the cell, and recordings were discarded when the Ca^2+^ current rundown exceeded 25%.

### Spinning disk confocal imaging of Ca^2+^ and iGluSnFR

Imaging experiments were performed with a spinning disk confocal scanner (CSU22, Yokogawa, Germany) mounted on an upright microscope (Axio Examiner, Zeiss, Germany) with 63x, 1.0 NA objective (W Plan-Apochromat, Zeiss). The spinning disk speed was set to 2000 rpm to avoid uneven illumination. A scientific CMOS camera (Neo, Andor, Northern Ireland) with a pixel size of 103 nm was used to acquire images. iGluSnFR and Rhod-FF or TAMRA-peptide were excited by diode-pumped solid-state lasers with 491 nm and 561 nm wavelength, respectively (Cobolt AB).

#### iGluSnFR imaging

To avoid photobleaching, iGluSnFR-expressing SGN boutons were detected via low intensity 491 nm excitation. The imaging plane for the target IHC was selected when several transduced boutons were visible, in the mid-basal section of the cell, avoiding the high synapse density at the basal pole. A brief step depolarization was applied to the cell to check for the functional signal in the given plane. iGluSnFR fluorescence was acquired at 50 Hz simultaneously with patch-clamp recordings. The iGluSnFR signal was evoked by step depolarizations of different durations to different voltage values, as it is specified in every dataset.

#### Sequential dual-color imaging of Ca^2+^ and iGluSnFR

For the sequential dual-color imaging of Ca^2+^ and glutamate release, as described above for iGluSnFR imaging, the imaging plane was selected based on the baseline fluorescence of iGluSnFR. Once the middle plane was set, the fluorescence of Rhod-FF was imaged at 100 Hz while Ca^2+^ currents were triggered by applying 5 voltage ramps (from −87 to +63 mV, 1 mV/ms) in 5 alternating planes separated by 0.5 μm. To precisely control the Z-plane, a piezo positioner for the objective (Piezosystem, Germany) was used. After the Ca^2+^ imaging, the iGluSnFR signal was acquired at 50 Hz by applying 50-ms-long step depolarizations from the holding potential of −87 to different voltage values. Depolarizations (to −57, −49, −45, −41, −37, −33, −25, −17 mV) were applied in a pseudo-randomized manner and covered the dynamic range of IHC glutamate release.

## Data Analysis

### Patch-clamp recordings

Electrophysiological recordings were analyzed using custom written programs in Igor Pro 6.3. Whole-cell Ca^2+^ charge (Q_Ca_) was calculated by the time-integral of the leak-subtracted current during the depolarization step. ΔC_m_ was calculated as the difference between the average C_m_ 400 ms before and after the depolarization, skipping the initial 100 ms after the depolarization.

### Imaging of iGluSnFR

Image and further data analysis and visualization were done in Python (Python software foundation) with custom written code using the following Python libraries: NumPy, Pandas, Matplotlib, Skimage, SciPy, Sklearn, statmodels and Seaborn.

#### Region of interest (ROI) detection

The ΔF image was created by subtracting baseline fluorescence (F_0_, an average of 15 frames before stimulus) from the fluorescence images acquired during/after stimulation (F, an average of 5 frames). The ΔF image was median-filtered with a size of 4-6 depending on the signal amplitude. To create a mask for ROI detection, maximum entropy thresholding was applied to the median-filtered ΔF image. To label and separate individual ROIs, a watershed segmentation algorithm was used. A single mask was generated per cell, using the recording with strongest stimulation, and applied for all images. Individual ROIs, corresponding to postsynaptic SGN boutons, were further confirmed by the nearby presence of presynaptic ribbon peptide (TAMRA-conjugated dimeric CtBP2-binding peptide). The fluorescence of every pixel in the defined ROI was averaged over time for further analysis. The background fluorescence was calculated by averaging 60 × 60 pixels in the pillar region of the image, where no iGluSnFR fluorescence is expected: By their anatomy, SGNs innervate IHCs and leave the cochlea towards the modiolus.

#### Analysis of fluorescence traces

The average background value was subtracted from the raw fluorescence traces (F). Following background subtraction, ΔF traces were generated by subtraction of mean baseline fluorescence (F_0_). ΔF was normalized to F_0_ to create ΔF/ F_0_ traces. For peak detection, ΔF/F_0_ traces were smoothened using a Hanning window function with a window size of 7. To correct for photobleaching, we fitted a single exponential to ΔF/F_0_ traces. The area under the curve (AUC) was estimated by calculating the area between ΔF/F_0_ and the fit in an interval of 40 frames from the beginning of the stimulus.

#### Estimation of sensitivity of iGluSnFR and ΔC_*m*_

For iGluSnFR, mean of 10 frames before stimulation is compared pairwise per synapse with the mean of 4 frames after stimulation. For ΔC_m_, mean of 400 points before and after stimulation is used for pairewise comparison per cell.

#### Estimation of the time to peak and decay constant

To obtain decay constant (τ_off_), the following function was fitted to the 30 points of photobleaching corrected ΔF/F_0_ traces after stimulation.

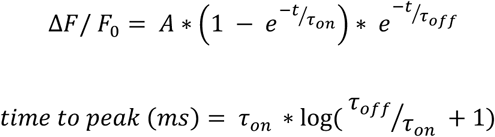

#### Estimation of the RRP size and time constant of RRP depletion

To assess the dynamics of RRP and sustained exoctosis, we fitted a sum of an exponential and linear function^22^ to ΔCm and iGluSnFR-AUC for different stimulus durations.

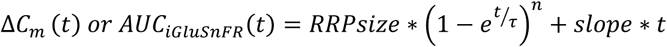

With the assumptions of ΔC_m_ of ∼40 aF contributed by a single SV^64^ and ∼12 AZs^28^ for the apical IHCs, we obtained iGluSnFR-AUC increase of 0.23 a.u. per SV.

### Sequential imaging of Ca^2+^ and iGluSnFR

#### ROI detection – iGluSnFR

The ROIs were picked as described above. Differently, a Gaussian filter with sigma of 1-3 was applied consistent with the detection of Ca^2+^ hotspots (see below). A mean mask was generated per cell using all the recordings. ROIs were confirmed by the presence of a corresponding Ca^2+^ “hotspot”.

#### ROI detection – Rhod-FF

Similarly, a ΔF image was created from the mean time series, in this case, by averaging all the trials from 5 recorded planes. This ΔF image was treated the same way as described for iGluSnFR-ROI picking. The created mask was applied to all recording planes and the plane with the maximum ΔF for a given ROI was used for further analysis. This way we used the plane with the highest signal for each Ca^2+^ “hotspot”.

#### Analysis of fluorescence traces

To remove the noise caused by the spinning disc speed at 2000 rpm (33.3 Hz), obvious in the Fourier amplitude spectrum, the raw traces were filtered with a 33.3 Hz bandstop filter. The obtained traces were background-subtracted and normalized to F_0_ as described above. Fluorescence-voltage (FV) relations for iGluSnFR were estimated from the step depolarizations to different voltage values. iGluSnFR-AUC for each depolarization was calculated as described above.

#### Estimation of the parameters of threshold, dynamic range, and V_1/2_

A Boltzmann fit was used to estimate the two fitting parameters: voltage of half-maximum activation (V_1/2_) and slope factor (*k*) of the glutamate release.

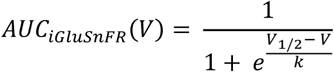

For Ca^2+^ imaging, FV curves were estimated from voltage ramps. To optimize the raw FV traces against noise such as readout or shot noise from the CCD camera, the following equation was used^11^:

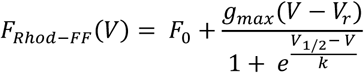

The slope factor (*k*) was obtained with this equation. The resulting fit was used to estimate V_1/2_ by minimizing the scalar at the mid-point. The reversal potential (V_r_) was fixed to +47.6 mV after LJ potential (17 mV) correction. In addition, this fit was used to calculate the fractional activation curve, dividing it by the extrapolated linear fit to the decay of fluorescence. To estimate fractional V_1/2,Pactivation_ and *k*_1/2,Pactivation_ (see Fig. 5), an additional Boltzmann fitting was done.

The peak of the Rhod-FF signal was obtained by averaging 3 frames corresponding to the voltage values between −17 mV and 3 mV during ramp depolarization. Dynamic ranges were calculated from the fits as the voltage range of 10%-90% of the maximal activation. Threshold was calculated as the voltage value where there is 10% of the maximal activation. Note that V_1/ 2_ values obtained from fluorescence traces after denoising (bandstop filter at 33.3 Hz) were comparable to the ones from the raw fluorescence traces.

#### Estimating the Ca^2+^ cooperativity

FV fits for Rhod-FF or Q_Ca_ and iGluSnFR-AUC were plotted against each other in the voltage range of −57 to −17 mV. To obtain single-synapse glutamate release-Ca^2+^ signal/whole-cell Ca^2+^ charge relationship, a power function was fitted:

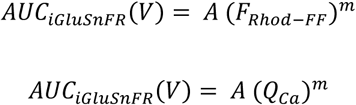

#### Calculation of the position of the synapse along cell’s pillar modiolar axis

To estimate the cell boundary, an ellipse was fitted to the baseline fluorescence of Rhod-FF. We defined the pillar-modiolar axis of the cell as the major axis of the ellipse. We calculated the shortest distance from the center of a given Ca^2+^ hotspot to the normalized major axis. A number was assigned to a given synapse on the normalized scale from 0 (pillar side) to 1 (modiolar side).

#### K-mean clustering and Principal Components Analysis

K-means clustering algorithm (K =3) was applied to the whole dataset of 11 synaptic properties for 3 clusters (see Fig. 6). Principal component analysis was used to display the clusters obtained by the K-means clustering.

### Statistical analysis

All the statistical tests were performed in Python (Python Software Foundation). Averages are expressed as mean ± SD, box plots indicate 25-75 quartile with whiskers reaching from 10-90%. Data sets were checked for normal distribution by D’Agostino & Pearson omnibus normality test and for equality of variances. For normally distributed data, unpaired two-tailed student’s t-test was applied, and for non-normally distributed data Mann-Whitney U test was used. Comparison of dispersion was performed by Levene’s test. We used one-way ANOVA for multiple comparisons followed by post-hoc Tukey’s test. The Pearson correlation coefficient was used to test for linear correlation. Significant differences are reported as * *p* ≤ 0.05, ** *p* ≤ 0.01, *** *p* ≤ 0.001.

## Supplementary Information

**Supplementary Figure 1:**
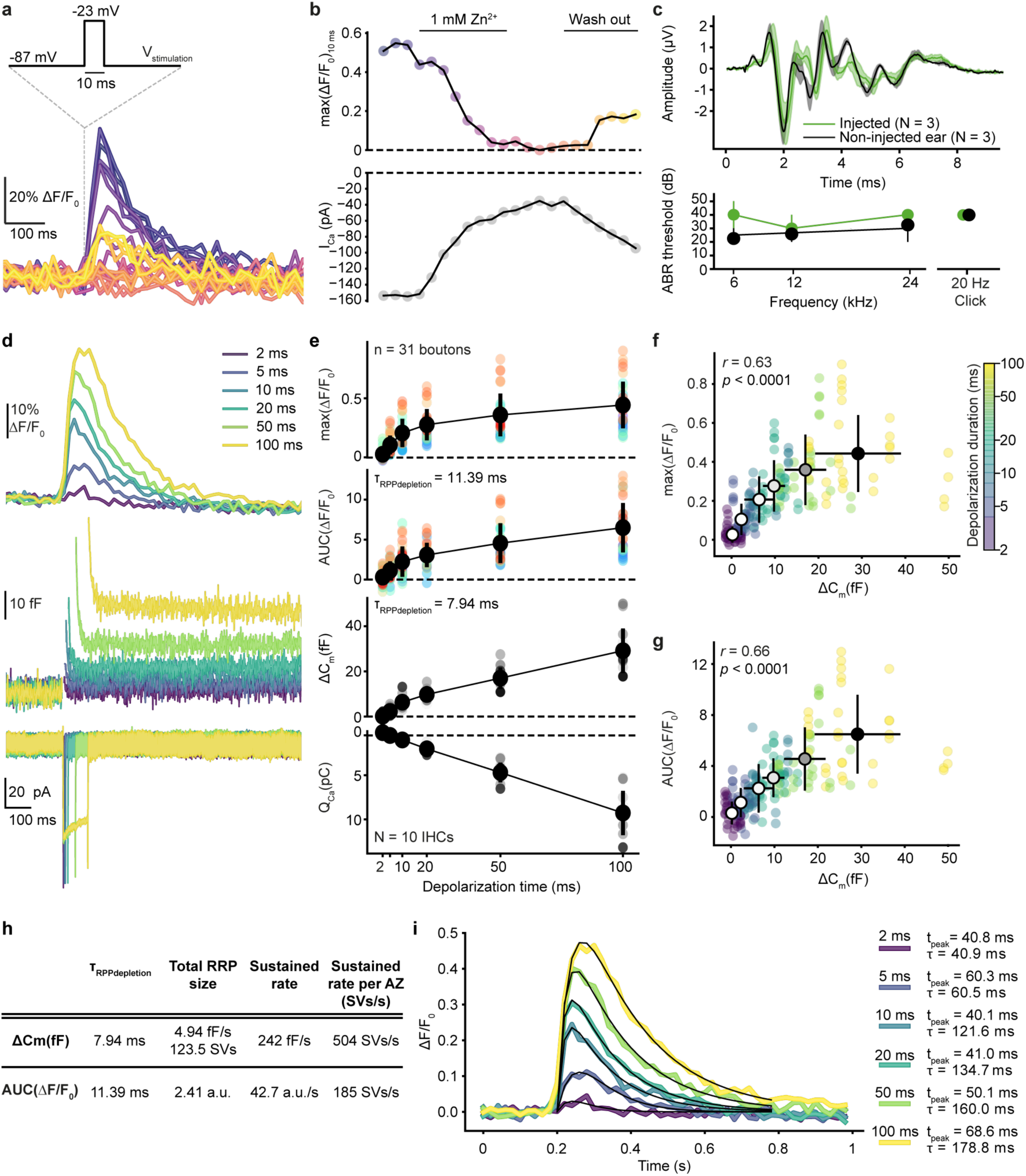
Characterization and validation of iGluSnFR for reporting IHC exocytosis. Related to Figure 1. **a.** Exemplary single-synapse iGluSnFR signal in response to repetitive 10-ms-long step depolarizations to −23 mV from a holding potential (−87 mV) as Ca^2+^ channel blocker Zn^2+^ (1 mM) is perfused in and out of the recording chamber. The temporal sequence of the recordings is encoded by color; darker colors indicate earlier time points as in panel B (top). **b.** The time course of the peak iGluSnFR response (top; max(ΔF/F_0_)_10ms_) from A and corresponding whole-cell peak Ca^2+^ current (bottom; perforated-patch, 1.3 mM [Ca^2+^]_e_). The whole-cell Ca^2+^ influx decreases with the perfusion of Zn^2+^. **c.** (top) ABR waveforms of P29 WT mice, injected with AAV9.*hSyn*.iGluSnFR virus at P6, were recorded in response to 80 dB clicks (mean ± SEM, 3 animals). The non-injected ear was used as a control. (bottom) ABR thresholds of the injected ear and the non-injected control were comparable. A statistical test was not applied due to small sample size. The presence of iGluSnFR expression was confirmed after ABR recordings. **d-g.** iGluSnFR signal as a readout of glutamate release increases with stimulus duration along with the IHC’s capacitance change (ΔC_m_). **d.** Average responses of iGluSnFR (top), whole-cell C_m_ (middle) and Ca^2+^ currents (bottom) upon step depolarizations to −23 mV from a holding potential of −87 mV for durations from 2 ms to 100 ms (color coded). Recordings were done in organs of Corti of P15-19 WT mice injected with AAV9.*hSyn*.iGluSnFR virus (perforated patch-clamp, 1.3 mM [Ca^2+^]_e_, *n* = 31 boutons, N = 10 IHCs from 8 mice). **e.** The peak and the AUC of iGluSnFR signal, corresponding whole-cell ΔC_m_, and Q_Ca_ plotted as a function of depolarization duration (mean ± SD). **f-g.** The relation of whole-cell ΔC_m_ and the peak (f) or the AUC (g) of the iGluSnFR signal (mean ± SD). Both the peak and the AUC of iGluSnFR response correlate with the whole-cell ΔC_m_ (Pearson’s *r* = 0.63, *p* <0.0001 and *r* = 0.66, *p* <0.0001, respectively). Depolarization duration is color coded, and the black outlined circles, indicating the means, darken with increasing depolarization duration. **h.** Quantification of exocytosis by whole-cell ΔC_m_ and single-synapse iGluSnFR-AUC. (See Methods). **I.** The kinetics of iGluSnFR signal. Average iGluSnFR responses (as shown in panel d top). Black lines indicate the results of fitting to the average traces per depolarization duration (see Methods). Time to peak and the decay constants are obtained from these fits and depicted with the color codes of the depolarization durations.

**Supplementary Figure 2.**
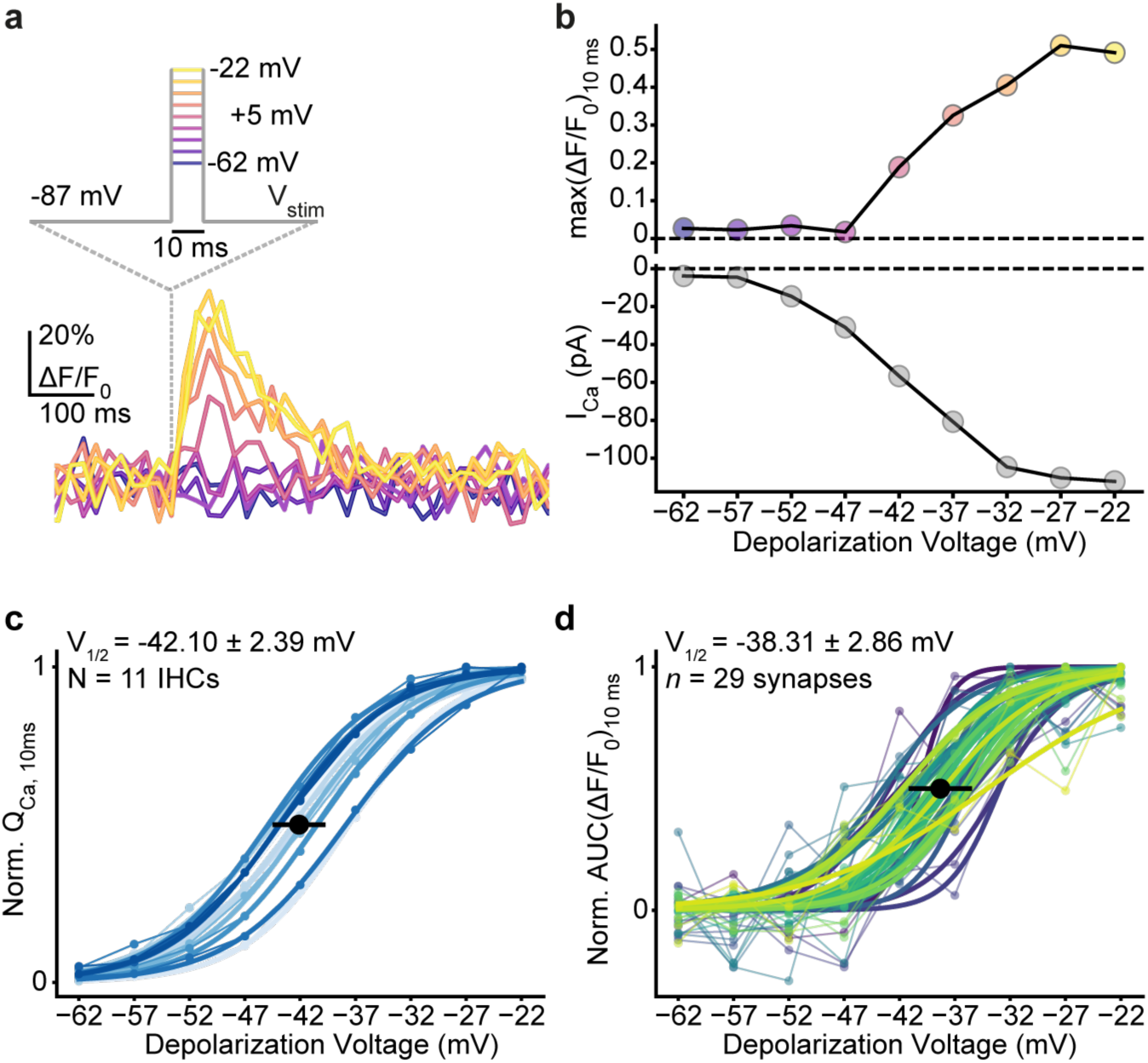
Low voltage threshold for IHC glutamate release. Related to Figure 1. **a.** Exemplary single-synapse iGluSnFR signal in response to 10-ms-long step depolarizations from the holding potential of −87 mV to a voltage within the physiologically relevant range of receptor potentials: from −22 mV to −62 mV in 5 mV increments, same as in Figure 1C-D. **b.** The peak iGluSnFR fluorescence (top; max(ΔF/F_0_)_10ms_) from A and corresponding whole-cell Ca^2+^ current (bottom; perforated patch-clamp, 1.3 mM [Ca^2+^]_e_). Glutamate release can be detected at −42 mV. The voltage range is color-coded: lighter points indicate more positive potentials. **c.** Normalized whole-cell Q_Ca_, calculated in response to 10-ms-long step depolarizations, is plotted as a function of depolarization voltage (perforated patch-clamp, 1.3 mM [Ca^2+^]_e_). Individual IHCs are color coded in shades of blue (mean ± SD, N = 11 IHCs from 9 mice). **d.** Normalized iGluSnFR AUC, in response to 10-ms-long step depolarizations, same experiments as in A (*n* = 29 synapses; individual synapses are color coded). A Boltzmann function was fitted to estimate the V_10_ and V_1/2._

**Supplementary Figure 3:**
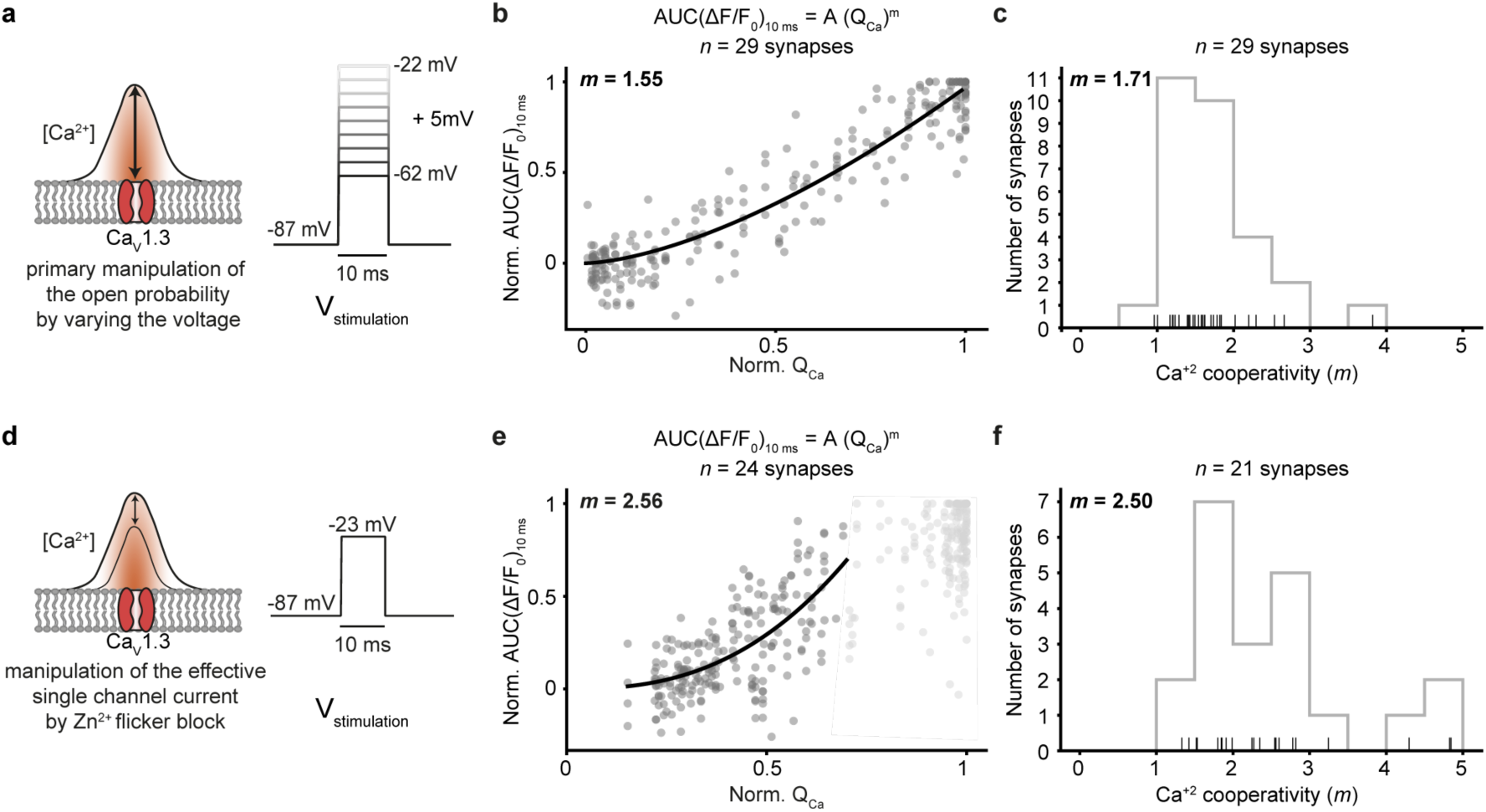
Relating release to whole-cell Ca^2+^ influx during manipulation of open-channel number or single-channel current supports Ca^2+^-nanodomain-like control of release. Related to Figure 1 and 3. **a.** Varying the voltage in the hyperpolarized range primarily varies the open-channel number. RRP release was probed by 10-ms-long step depolarizations from the holding potential (−87 mV) to −62 mV to −22 mV in 5 mV steps applied in pseudo-randomized order. **b.** The normalized AUC(ΔF/F_0_)_10ms_ is plotted versus Q_Ca_ (*n* = 29 boutons, N = 11 IHCs from 9 mice): a power function was fitted before an obvious saturation of the RRP release, and a near-linear relation was observed (*m* = 1.55). **c.** Histogram showing the distribution of *m* from individual fits per synapse before an obvious saturation of the RRP release (*m*_*avg*_ = 1.71 ± 0.58). Only those fits with an R^2^ value higher than 0.7 were used for further analysis (*n* = 29 boutons). The rug plot under the histogram displays individual data points. Every depolarization step was repeated at least two times per synapse and the average was taken. **d.** Manipulation of the single Ca^2+^ channel current by Zn^2+^ perfusion. Note that Zn^2+^ acts as a rapid (microsecond) flicker blocker of Ca^2+^ channels^1^, which, within the limits of IHC exocytosis kinetics (delay ∼1 ms)^2^ is expected to reduce the fusogenic Ca^2+^ signal at the SV release site^3^. Therefore, this manipulation is expected to reveal the supralinear intrinsic Ca^2+^ dependence of release. We evoked RRP exocytosis by repetitive 10-ms-long step depolarizations to −22 mV, while slowly perfusing 1 mM Zn^2+^ into the recording chamber. **e.** Normalized AUC(ΔF/F_0_)_10ms_ is plotted versus Q_Ca_ upon Zn^2+^ manipulation (*n* = 24 boutons, N = 10 IHCs from 10 mice). A power function was fitted before an obvious saturation of RRP release (normalized Q_Ca_ < 0.7), and a supralinear relation was observed (*m* = 2.56). **f.** Histogram showing the distribution of *m* from individual fits synapse before an obvious saturation was observed for the given synapse (*m*_*average*_ = 2.50 ± 1.03, *n* = 21 boutons with an R^2^ of fit > 0.7). The rug plot under the histogram displays the individual data points.

**Supplementary Figure 4:**
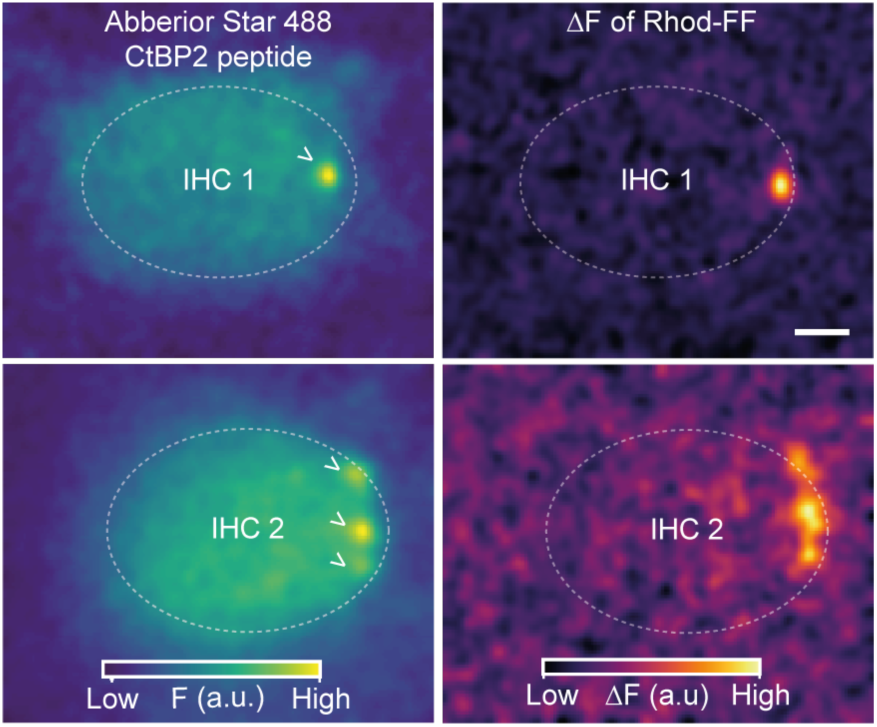
Depolarization-evoked hotspots of Rhod-FF fluorescence occur at the AZ. Related to Figure 2. **(Left)** Confocal sections of two example IHCs showing the fluorescence of the Abberior Star 488-conjugated CtBP2-binding peptide (the synaptic ribbons are marked with an arrowhead). **(Right)** ΔF images of Rhod-FF in response to 100-ms-long step depolarization to −17 mV from a holding potential of −87 mV. The color maps of the F and ΔF images in arbitrary units (a.u.) and are displayed on the bottom.

**Supplementary Figure 5:**
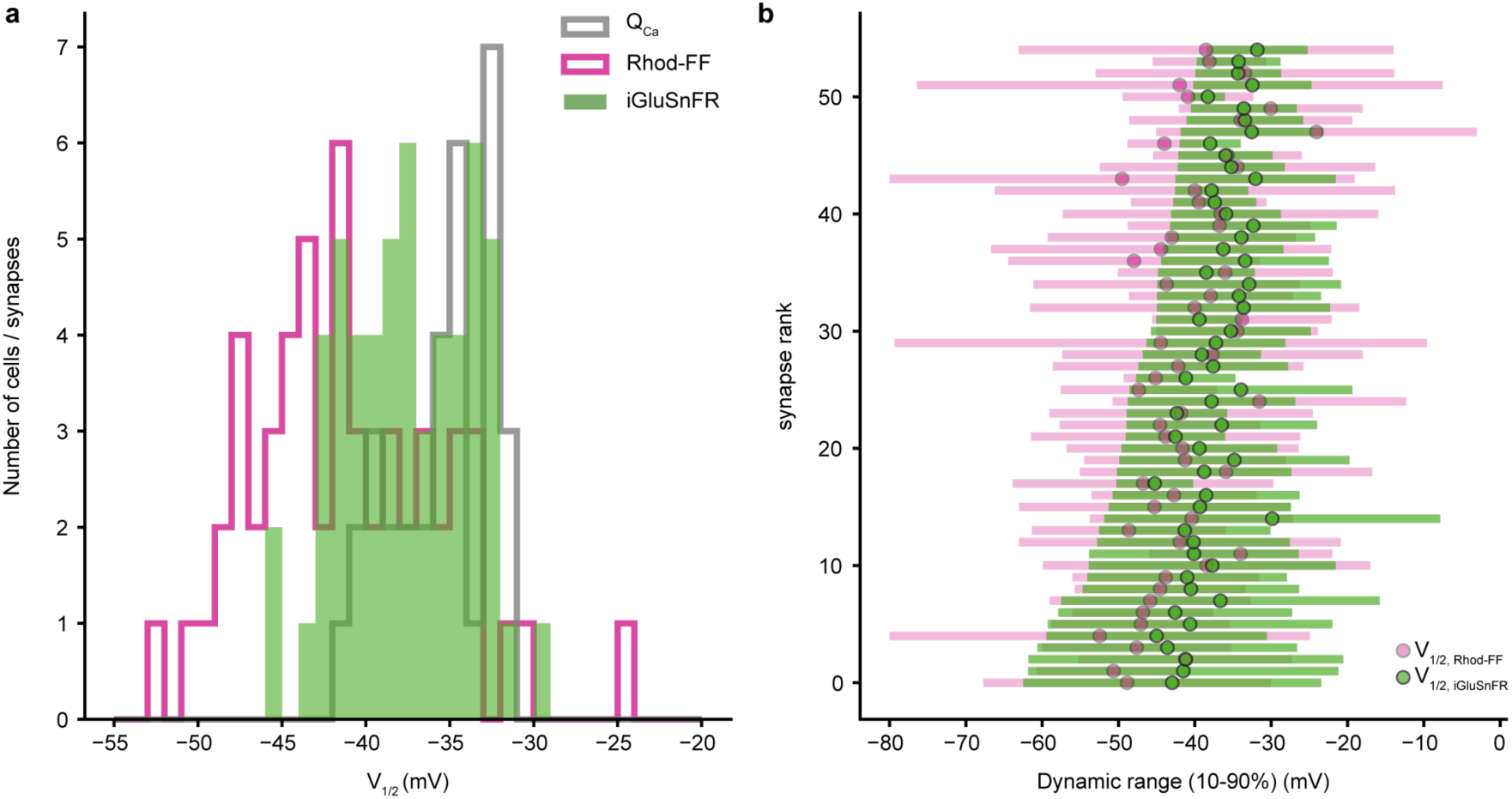
AZs vary in their voltage-dependence. Related to Figure 3. **a.** V_1/2_ distribution of Q_Ca_ (gray, N = 34 IHCs), synaptic Ca^2+^ influx (orange) and glutamate release (green, *n* = 55 synapses from 34 IHCs). **b.** Dynamic ranges of the synaptic Ca^2+^ influx (orange) and glutamate release (green) with their V_1/2_ depicted. The synapses are ranked based on their glutamate release threshold (V_10_). Note how they gradually span the voltage range from −62.48 mV to −38.41 mV (mean ± SD = 48.27 ± 6.47 mV).

**Supplementary Figure 6.**
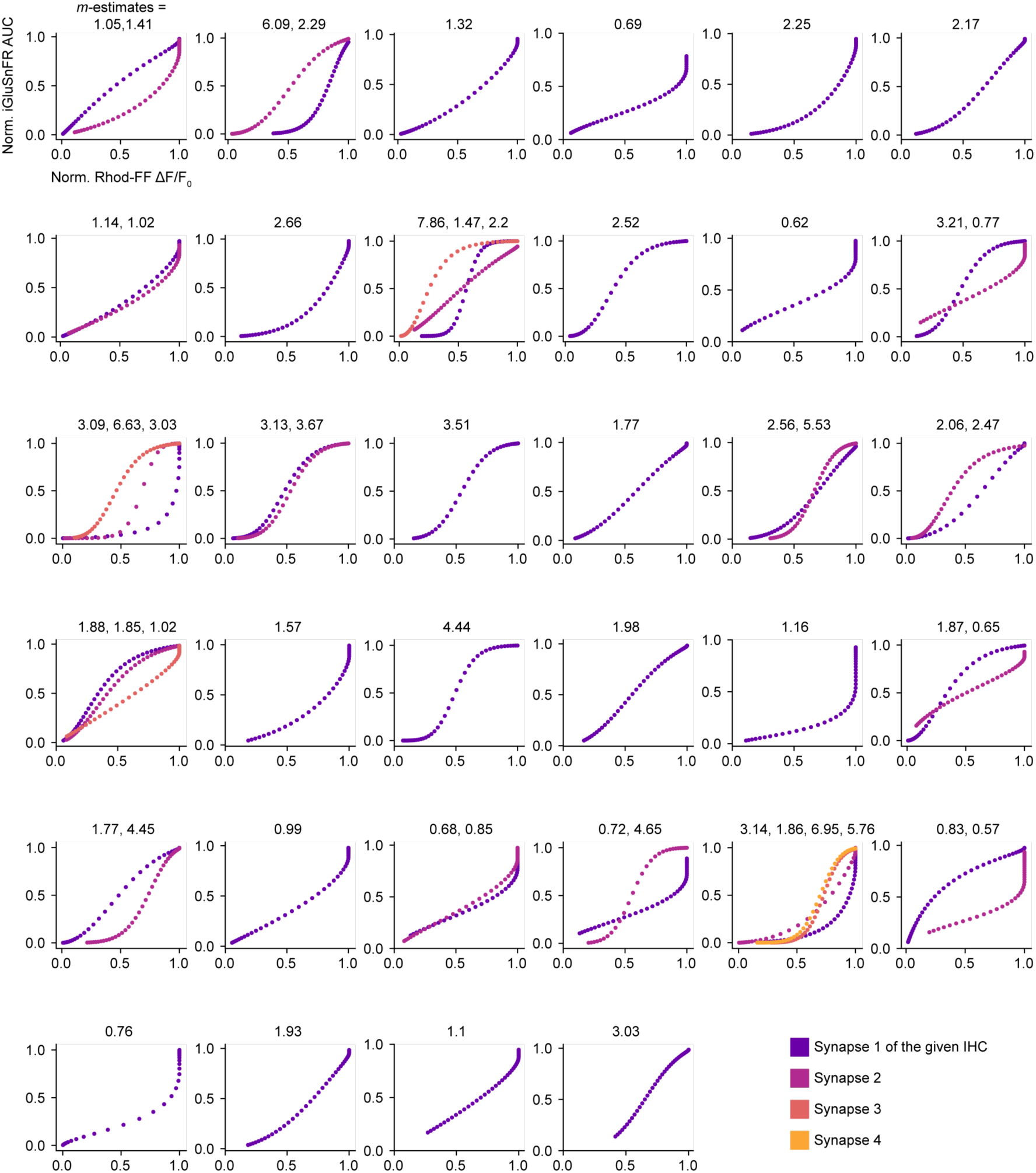
Apparent Ca^2+^ dependence of synapses in individual IHCs. Related to Figure 3. The relations of synaptic Ca^2+^ influx (normalized modified Boltzmann fit on ΔF/F_0_ of Rhod-FF) and glutamate release (normalized Boltzmann fit on iGluSnFR-AUC) were plotted per IHC. Ca^2+^ cooperativities (*m*-estimates) were obtained by individual power function fitting until 25% of normalized iGluSnFR response. Synapses of a given IHC are color coded and displayed on the lower right corner.

